# Obesity restricts oligodendrocyte maturation and impedes repair after white matter stroke

**DOI:** 10.1101/283184

**Authors:** Guanxi Xiao, Jasmine Burguet, Riki Kawaguchi, Leif A. Havton, Jason D. Hinman

## Abstract

Obesity is a growing public health problem that increases rates of white matter atrophy and increases the likelihood of ischemic lesions within white matter. However, the cellular and molecular mechanisms that regulate these changes are unknown. We hypothesized that obesity may alter oligodendrocytes and myelin priming white matter for worsening injury and repair responses after ischemia. C57Bl/6 mice fed a high fat diet (60% kcal from fat) show increased numbers of oligodendrocyte progenitor cells (OPCs), decreased myelin thickness with elevated g-ratios, and shorter paranodal axonal segments, indicating compromised myelination in high fat diet mice. Fate mapping of OPCs in *PDGFRα-Cre^ERT^;Rpl22^tm1.1Psam^* mice demonstrated that OPC differentiation rates are enhanced by obesity. Gene expression analyses using a novel oligodendrocyte staging assay demonstrated that obesity restricts oligodendrocyte maturation in between the pre-myelinating and myelinating stages. Using a model of subcortical white matter stroke, the number of stroke-responsive OPCs in obese mice was increased after stroke. At early time points after ischemic white matter stroke, spatial mapping of stroke-responsive OPCs indicates that obesity leads to increased OPCs at the edge of ischemic white matter lesions. At later time points, obesity results in increased OPCs within the ischemic lesion while reducing the number of GST-π-positive mature oligodendrocytes in the lesion core. These data indicate that obesity disrupts normal white matter biology by impeding oligodendrocyte differentiation, leading to an exaggerated response of OPCs to white matter ischemia, and limited brain repair after stroke.

---

Main points
- Adult-onset obesity reduces myelin thickness, increases oligodendrocyte precursor cell (OPC) differentiation, and blocks oligodendrocyte maturation.
- After focal white matter stroke in obese mice, the early OPC response to stroke is exaggerated while late reparative OPC differentiation is impaired.

## Introduction

Obesity is an emerging public health crisis with rates of adult obesity in the US approaching 40% (Ogden et al., 2015). Among the various ill-health effects of obesity, its role in damaging brain white matter is clear. Obesity increases the risk of developing silent lacunar infarcts (Bokura et al., 2008; Park et al., 2008) six-fold and increases the detection of white matter hyperintensities on MRI (Park et al., 2008). Increased body mass index is also associated with white matter atrophy (Karlsson et al., 2013) and reduced white matter tract integrity (Stanek et al., 2011; Bettcher et al., 2013; Xu et al., 2013; Kullmann et al., 2016). These observed declines in fractional anisotropy (FA) using diffusion tensor imaging suggest that obesity specifically disrupts myelination and axonal integrity. These obesity-related white matter changes are increasingly linked with cognitive impairment (Debette et al., 2011) and functional disability (Inzitari et al., 2009).

Despite the increasing rates of obesity and its established relationship with white matter pathologies in humans, the effect of obesity on white matter biology remains largely unknown. In leptin-deficient genetically obese mice (ob/ob), lower amounts of myelin and the fatty acid composition of myelin are altered (Sena et al., 1985). At early embryonic stages, leptin-deficiency contributes to an increased number of oligodendrocyte progenitor cells (OPCs) (Udagawa et al., 2006), while leptin-treated ob/ob mice showed an increase in myelination (Gouw et al., 2008) suggesting that leptin may partially regulate oligodendrocyte differentiation. Whether these findings in leptin-deficient mice relate to the more widespread forms and models of diet-induced obesity is unclear.

Increased in frequency by obesity, focal ischemic white matter strokes are common (Benjamin et al., 2018) and progressive (Gouw et al., 2008). The pathology of these lesions revolves centrally around the oligodendrocyte lineage. Mature oligodendrocytes are exquisitely sensitive to ischemia (Oka et al., 1993; Pantoni et al., 1996; McDonald et al., 1998b; McDonald et al., 1998a), undergo rapid (Sozmen et al., 2009) and progressive (Hinman and Carmichael, 2014) cell death after focal white matter lesions. OPCs respond robustly to ischemic white matter lesions (Sozmen et al., 2016). Limited OPC differentiation and remyelination after white matter stroke tempers functional recovery, particularly in aged animals (Rosenzweig and Carmichael, 2013). Understanding the role obesity plays in normal white matter biology and its role in altering white matter injury and repair seems a pre-requisite to designing therapeutic strategies for this common form of stroke.

To examine the role obesity plays in altering white matter and the response to a focal white matter stroke, we used cellular, ultrastructural, and gene expression approaches in combination with diet-induced obesity and a mouse model of focal white matter stroke. We show that obesity increases the resting number of OPCs in white matter yet also blocks their maturation specifically at the transition point from pre-myelinating to mature myelinating oligodendrocytes leading to thinner myelin sheaths and axonal microdomain changes. In turn, these obesity-induced effects on the oligodendrocyte lineage result in an exaggerated response to white matter stroke creating a larger lesion burden and impaired OPC differentiation in response to stroke.

## Material & Methods

### Mice

All animal studies presented here were approved by the UCLA Animal Research Committee, accredited by the AAALAC. Mice were housed under UCLA regulation with a 12 hour dark-light cycle. All mice used in the study were male. Wild-type C57Bl/6 mice fed ad lib on 60%kCal from fat chow (HFD) (Strain #380050) or 10%kCal from fat chow (CFD) (Strain #380056) were purchased directly from Jackson Labs at 17 weeks of age and allowed to acclimate for 2 weeks prior to experimental use. The PDGFRα-CreERT^2^/Rpl22-HA transgenic strain was generated by crossing PDGFRα-CreERT^2^ mice (Jackson Labs Strain #018280 - B6N.Cg-Tg(Pdgfra-cre/ERT)467Dbe/J) with Rpl22-fl-Rpl22-HA (Jackson Labs Strain #011029 - B6N.129-*Rpl22^tm1.1Psam^*/J). Diet-induced obesity was induced in transgenic mice by ad lib feeding with 60%kCal from fat chow (HFD) or 10%kCal from fat chow (CFD) (Research Diets, Inc.). Weights (g) were measured weekly.

### Fate mapping

Tamoxifen (Sigma) were dissolved in corn oil to make 20mg/ml stock solution. For early stage OPC fate mapping, tamoxifen was injected i.p. to PDGFRα-CreERT^2^/Rpl22-HA (*n* = 9) mice (50mg/kg) daily for 4 consecutive days beginning at 8 weeks of age. Mice were euthanized and brains were analyzed at 20 weeks of age. For late stage OPC fate mapping, tamoxifen was injected i.p. to PDGFRα-CreERT^2^/Rpl22-HA (*n* = 8) mice (50mg/kg) daily for 4 consecutive day at 20 weeks of age. Mice were euthanized and brains analyzed at 28 days after the last injection.

### Electron microscopy

Wild-type C57Bl/6 mice (*n* = 6/grp) on CFD or HFD were transcardially perfused with a 2% glutaraldehyde solution, post-fixed for 24 hrs, hemisected in the sagittal plane and 2 mm cubes including the corpus callosum were dissected and embedded in plastic resin for ultrastructural analysis as previously described (Sozmen et al., 2016). One micron, plastic embedded toluidine blue stained sections were used to select transcallosal fibers underneath sensorimotor cortex by light microscopy. Three electron micrographs were obtained at a primary magnification of 7200X using a JEOL 100 CX transmission electron microscopeand a representative electron micrograph of high technical quality from each animal was used for quantitation of fiber diameter, axon diameter, myelin thickness, and g-ratio.

### Immunofluorescence

Animals were euthanized with a lethal dose of isoflurane, transcardially perfused with PBS followed by 4% paraformaldehyde in 0.1 M sodium phosphate buffer, brains removed, post-fixed for 24 hrs and cryoprotected for 48 hrs in 30% sucrose in PBS. Forty micron coronal cryosections and immunostaining was performed essentially as described (Hinman et al., 2013). The following primary antibodies were used: mouse anti-NF200 (1:200, Sigma), rabbit anti-MBP (1:500, Calbiochem), goat anti-PDGFRα (1:500; Neuromics), mouse anti-HA (1:1000, Biolegend), rabbit anti-AnkG (1:1000, Dako), rabbit anti-Na_V_1.6 (1:250), rabbit-Gst-π (1:1000, Millipore), in PBS containing 5% goat or donkey serum and 0.3% Triton-X 100 (Sigma) overnight at 4°C. Secondary antibody labeling was performed using donkey anti-mouse, donkey anti-rabbit, or donkey anti-goat Fab_2_-Alexa conjugated antibodies (Jackson Immunoresearch, Inc.).

### Microscopy

All microscopic images were obtained using a Nikon C2 confocal microscope. Optical slices were analyzed in Imaris with automated “Add Spots” function to generate cell counts and x,y,z positional coordinates. OPCs stroke responsive areas were generated by Imaris with “Add Surface” function. Measurements of stroke areas were generated using Fiji (Schindelin et al., 2012). Representative images were selected for presentation.

### Gene expression analysis

Using established oligodendrocyte stage marker genes (Zhang et al., 2014), we designed a custom Nanostring^®^ gene expression array (XT_GX CodeSet Oligostg #116000651) using gene-specific probes for each the 40 genes marking each oligodendrocyte stage (OPC, pre-myelinating oligodendrocyte (PMO), and myelinating oligodendrocyte (MO)). C57Bl/6 mice (n=4/group) were maintained on 12 weeks of CFD or HFD. At 20 weeks of age, the corpus callosum was isolated, homogenized, and RNA using the Nucleospin miRNA kit (Clontech) to collect both large and small RNA species. RNA was quantitated and 100 ng of RNA from each animal was provided as input for the direct RNA detection assay. Manufacturer protocol was followed and the results were analyzed using nCounter^®^ software. Results were normalized and differentially expressed genes were compared individually between groups using a Student’s t-test (*p*<0.05). The number of DEGs (*p*<0.05) per stage was determined. log2FC values were calculated as a function of averaged housekeeping gene expression levels (*actb, b2m, gapdh, pgk1, rpl19*). To generate an oligodendrocyte stage cell type index, raw gene expression values were combined with FPKM values from Zhang et al. (2014) and normalized by rank. Hierarchal clustering analysis was performed using hclust and tSNE plots (perplexity=3, max iteration = 5000) generated using standard algorithms. The resulting classification and 2-dimensional representation were highly reproducible.

### White matter stroke

Subcortical white matter ischemic injury was induced as previously described (Hinman et al., 2013; Nunez et al., 2016). Animals (*n* = 4/grp) were sacrificed at 7 and 28 days post-stroke and analyzed for tissue outcomes.

### Spatial analysis

Analysis of the spatial distribution of stroke-responsive OPCs was performed as follows. The boundary of increased PDGFR-α-+ cells and the loss of GST-π-+ cells was identified in each of three sections per animal (*n*=3 animals/group). Using Imaris software, the x,y,z position of each PDGFR-α-+ cell relative to the user defined center point (x=0, y=0, z=0) of the elliptical stroke region was determined. Because the z-axis was limited (10 μm), a two-dimensional grid analysis was performed using a 2D modification of the previously reported 3D spatial density estimator (Burguet et al., 2011) using a smoothing parameter of *k*=8. The local cell density in each position within the overlaid grid is compared statistically as previously described (Burguet and Andrey, 2014). Therefore, a *p-*value map is generated for each position in the grid and thresholded (*p*<0.05) to reveal regions with significant density differences.

### Experimental Design and Statistical Analysis

The number of animals used in each experiment is listed in the Results section. Oligodendrocyte population cell counts as a fraction of total cells were determined by averaging counts from 5 fields of view throughout the corpus callosum across a minimum of three sections 240 μm apart. Per animal averages were generated and significance between groups determined using an unpaired Welch’s t-test (α=0.05). For OPC fate mapping, similarly obtained per animal averages were analyzed using Holm-Sidak test to adjust *p-*values for multiple t-tests. Analysis of nodal/paranodal complexes was performed using an unpaired Welch’s t-test (α=0.05), while paranodal length varies based on axonal diameter and was therefore analyzed using a distribution analysis and Chi-square statistic (*df*=12). Measurements of white matter ultrastructural features were averaged across animals and each feature was compared separately using Mann-Whitney U test between groups (α=0.05). Gene expression differences were determined at the individual gene level using unpaired Welch’s t-test (α=0.05). Determination of stroke lesion area was performed by sampling lesion area (*n*=3–5 40 μm sections) across groups (*n*=4/grp) and using the sampled distribution to create bootstrapped area distribution (*n*=25) representing a full area sampling of the approximate 1 mm lesion created by the stroke model. This area distribution was averaged across animals in each group and compared using a Mann-Whitney U test between groups (α=0.05). Spatial analysis of stroke-responsive OPCs were determined as detailed above. Cell counts at 28d post-stroke were determined across three sections 240 μm apart with lesion core and edge analyses determined using a two-way ANOVA (α=0.05) with post-hoc Holm-Sidak test to adjust *p-*values for multiple t-tests. Statistical analysis was performed using GraphPad Prism 7 software. Data are shown as mean ± SEM.

## Results

### Obesity increases the oligodendrocyte progenitor cell population in white matter

To induce obesity, mice were fed with 60% kCal from fat diet (HFD) beginning at 8 weeks of age and continuing for 12 weeks (Surwit et al., 1988; Podrini et al., 2013). Mice fed with HFD develop severe obesity with an average 84.0% weight gain compared to 21.7% in control diet fed mice (10% kCal from fat; CFD) (Fig. 1a). This weight gain is associated elevations of total cholesterol (110.7 vs. 187.3; *p*=0.023), LDL (37.0 vs. 99.7; *p*=0.043), glucose (162.0 vs. 373.3) and HgbA1c (5.6 vs. 6.6), largely consistent with the clinical definition of metabolic syndrome (Benjamin et al., 2018). To determine the effects of obesity on oligodendrocytes, we immunolabeled brain sections with PDGFRα to label OPCs and GST-π to label mature oligodendrocytes (Fig. 1c-f). We sampled oligodendrocyte cell populations throughout the frontal subcortical white matter (Fig. 1b). The percentage of GST-π+ cells were nearly identical in CFD (76.4±1.4%) and HFD (75.8±0.6%) mice (Fig. 1g). However, the percentage of PDGFRα+ cells was significantly increased in HFD (5.66±0.22%) compared to CFD (5.01±0.13%; *p*=0.014; Fig. 1h). Importantly, PDGFRα+ cell counts in CFD animals were remarkably similar to previous reports of the abundance of the OPC population in white matter (Pringle and Richardson, 1993; Chang et al., 2000; Dawson et al., 2003; Bergles and Richardson, 2015). These data indicate that obesity modestly elevates the resting proportion of OPCs in white matter.

**Figure 1.**
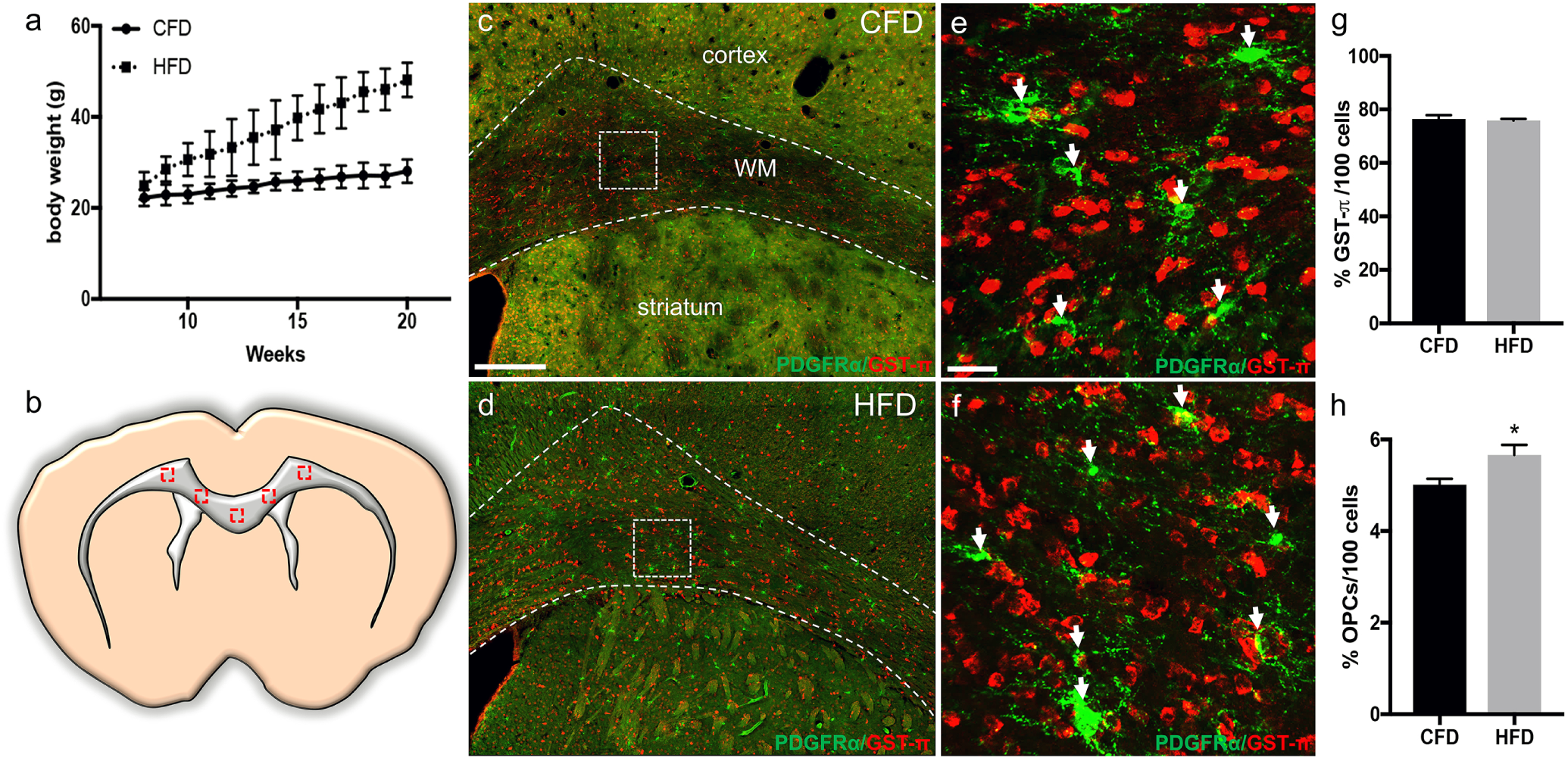
High fat diet increases oligodendrocyte precursor cells in white matter. Consumption of a 60% kcal from fat diet (HFD) results in rapid and significant weight gain beginning from week 9 compared to 10% kcal from fat diet (CFD) (a). After 12 weeks on HFD, we sampled corpus callosum and subcortical white matter as indicated (red boxes) (b). Labeling for GST-π+ mature oligodendrocytes (red) and PDGFR-α+ OPCs (green) in animals on CFD (c, e) and HFD (d, f) reveals an increase in PDGFR-α+ cells (arrows, e vs. f). Quantitation of GST-π+ cells/100 cells was not changed (g) while the percentage of PDGFR-α+ cells was significantly increased in animals on HFD (h). Data are mean ± SEM. Scale bars = 200 μm (c, d) and 20 μm (e, f).

### Obesity compromises axonal integrity in white matter

Because obesity increased the OPC population in white matter, we reasoned that myelin turnover might be affected leading to changes at nodal and paranodal structures within axons. We examined axonal microdomain structure in callosal axons using immunolabeling for Na_V_1.6 to label the node and Caspr to label the paranodal segment (Fig. 2a-b). In mice on CFD, the nodal and paranodal structures were normal (Fig. 2a inset) while in mice on HFD, paranodal segments appeared shorter and the number of intact nodal and paranodal complexes were reduced (Fig. 2b) In HFD mice, we found a decrease in intact nodal (green)/paranodal (red) complexes (defined as node with two adjacent paranodes) per field (Fig. 2c). Measurement of paranodal length was performed and the frequency of paranodes within each 0.2 µm bin from 0.6–3.0 µm in length were determined. In mice on HFD, we found a reduction in average paranodal length from 1.70±0.03 µm in CFD (*n*=256 paranodes) compared to 1.29±0.02 µm (*n*=337 paranodes; *p*<0.0001) which was associated with a significant left shift of the distribution of paranodal lengths (*x^2^*=1×10^−137^) (Fig. 2d), indicating shorter paranodal segments consistent with incomplete myelination.

**Figure 2.**
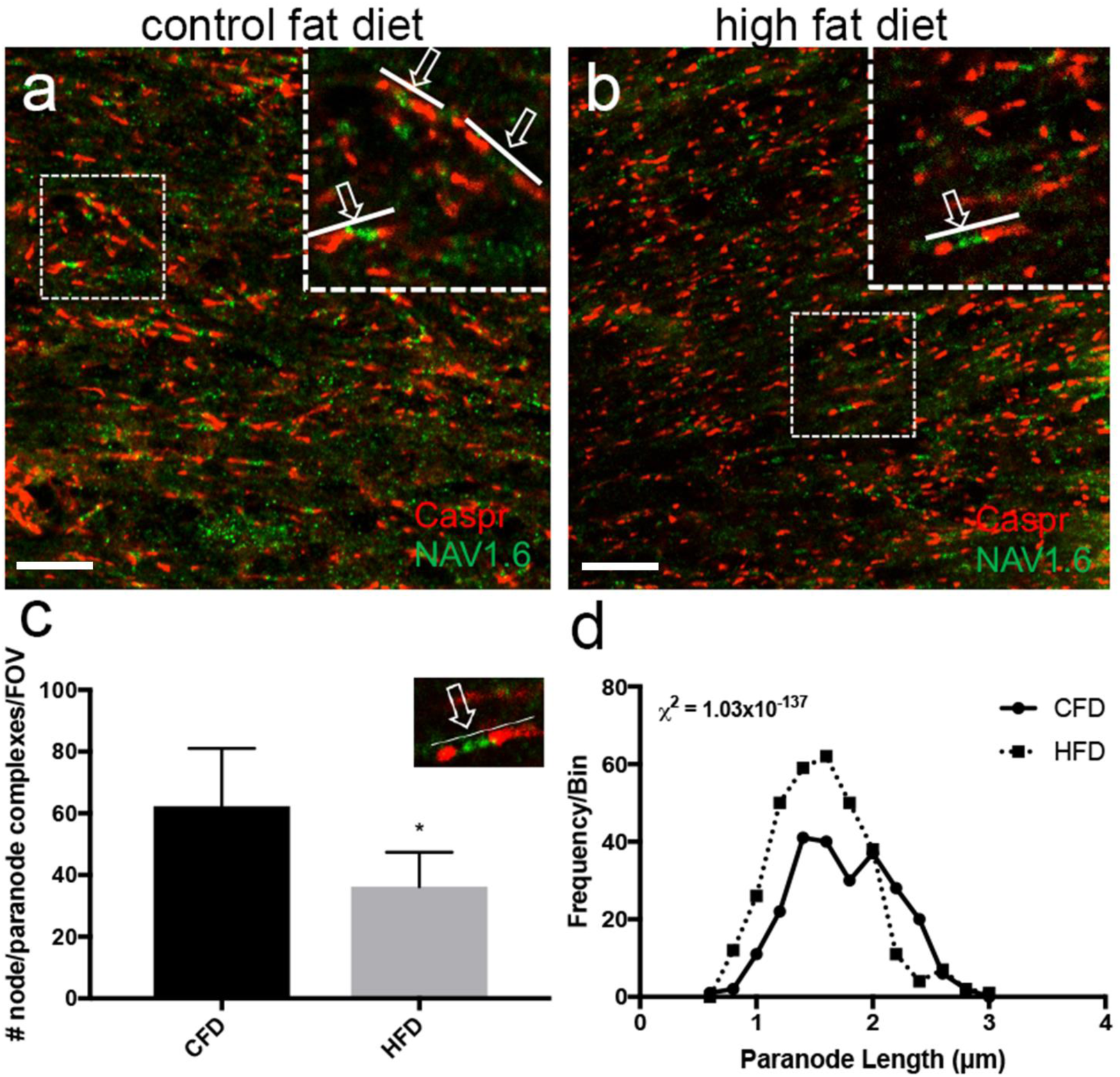
High fat diet reduces axonal microdomain integrity and paranodal length. Labeling for nodal (Na_V_1.6, green) and paranodal (caspr, red) axonal microdomain segments within white matter in animals on CFD (a) and HFD (b). Inset boxes show examples of intact nodal and paranodal complexes (arrow with line). The number of nodal and paranodal complexes, defined as adjacent caspr+ paranodal segments with concurrent Na_V_1.6+ nodal segments (example in inset), is reduced in animals on HFD (c). The distribution of paranodal length (binned by 0.2 μm) demonstrates a significant right shift towards shorter paranodes in animals on HFD (dashed line) compared to CFD (solid line) (d). Scale bar = 10 μm.

### Obesity reduces myelin thickness

To firmly establish the effects of obesity on myelin and white matter, we prepared sagittal sections including callosal fibers for ultrastructural analysis by EM. In 20-week old mice on CFD (*n* = 6), myelin ultrastructure was quite normal with intact axonal fibers, myelin thickness, and a g-ratio of 0.8 (Fig. 3a). In 20-week old mice on HFD (*n* = 6), we observed an increased frequency of moderate to larger size axonal fibers with thinner than expected myelin sheaths (Fig. 3b). The total number of axonal fibers measured was not significantly different between CFD (135.8±10.48) vs. HFD (107.2±14.88; *p*=0.18). Neither the average fiber diameter (0.76±0.07 μm vs. 0.79±0.06 μm; *p*=0.81) nor axonal diameter (0.62±0.06 μm vs. 0.69±0.05 μm; *p*=0.48) were significantly different between mice on CFD vs. HFD. Average myelin sheath thickness was significantly reduced in mice on HFD (Fig. 3c; 0.070±0.005 μm vs. 0.047±0.004 μm; *p*=0.0087) and the average g-ratio in animals on HFD was consequently increased compared to those on CFD (Fig. 3d; 0.8±0.011 vs. 0.88±0.006; *p*=0.002). Plotting individual axon diameter vs. g-ratios for all measured axons by dietary status reveals a shift towards higher g-ratios in small and larger caliber axons (Fig. 3e) in mice on HFD (red) compared to those on CFD (black). The Pearson correlation coefficients for g-ratios and axon diameter were substantially different between CFD (0.67) and HFD (0.45). These results indicate that obesity leads to thinner myelin sheaths within the corpus callosum.

**Figure 3.**
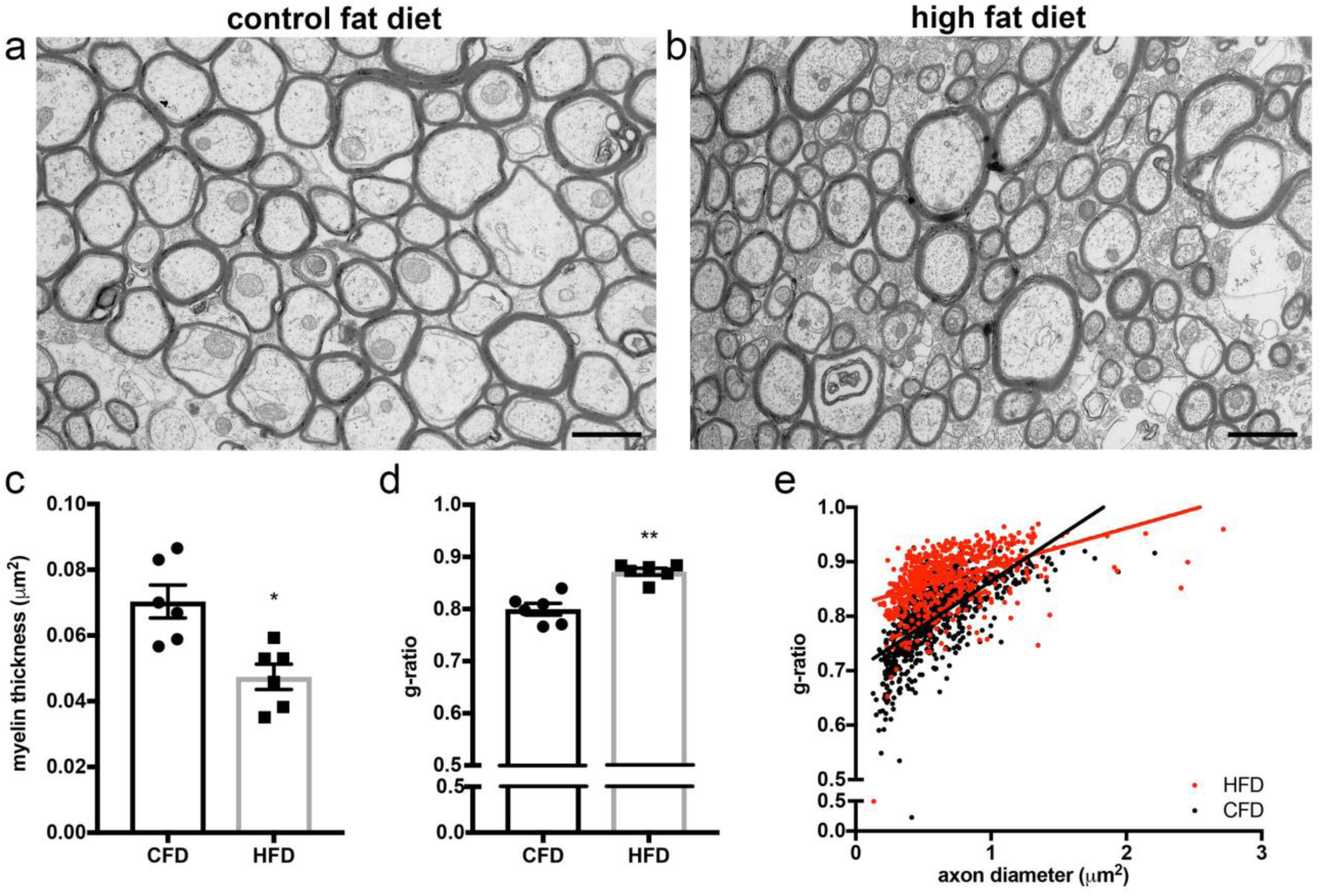
High fat diet compromises myelin ultrastructure. Callosal fibers at the midline were prepared in sagittal section for electron microscopy. Animals on CFD demonstrated regular axonal and fiber diameter with an abundance of normally myelinated fibers (a). After 12 weeks on HFD, myelin ultrastructure in callosal fibers is compromised with reductions in myelin sheath thickness without significant changes in axonal number or fiber diameter (b). Myelin sheath thickness was significantly reduced (0.070 μm vs. 0.047 μm; *p*=0.0087) (c). The average g-ratio was increased (0.88 vs. 0.80; *p*=0.002; *n*=6) in animals on HFD compared to CFD (d). The axon diameters were not significantly different between CFD and HFD (e).7200X magnification; scale bar = 1 μm. Data are mean ± SEM.

### Obesity accelerates oligodendrocyte differentiation

The increased proportion of OPC and compromised integrity of myelin and white matter resulting from obesity suggests a failure of OPC differentiation into myelin-producing mature oligodendrocytes. However, it is unclear how OPCs responses during the development of obesity and after obesity is present. To answer this question, we performed fate mapping at two different points to analyze OPC differentiation using a novel *PDGFRα-Cre^ERT^;Rpl22^tm1.1Psam^* transgenic mouse strain we created by crossing the *PDGFRα-Cre^ERT^* strain with the *Rpl22^tm1.1Psam^-*HA tagged ribosome reporter mouse (Fig. 4a). To confirm phenotypic labeling of OPCs, mice (*n*=2) were injected with tamoxifen and analyzed 48 hours after the injection. HA+ reporter cells were clearly co-labeled with PDGFRα, indicating tamoxifen administration clearly labels OPCs (Fig. 4b). We performed OPC fate mapping both before and after the onset of obesity. First, we analyzed the response of OPCs after obesity is developed. Tamoxifen was injected at 20 weeks in CFD (*n*=4) and HFD (*n*=4) mice (31.3±1.5g vs. 45.5±3.8g; *p*=0.014). The numbers of PDGFRα+/HA+ (6.5±0.7%) and HA+ (26.7±6.5%) cells were not significantly changed in HFD white matter compared to CFD (7.1±0.8%; 30.2±1.8% respectively; *p*=0.70) (data not shown). However, administration of tamoxifen prior to the onset of obesity allowed us to track the fate of OPCs during the development of obesity. Early fate mapping of OPCs labeled at 8 weeks of age concurrent with starting the HFD demonstrated a significant increase in HA+ cells in HFD white matter (68.8±1.60, *n*=4) compared to CFD (54.5±2.98; *n*=4; *p*=0.011), while the number of PDGFRα+/HA+ cells was not significantly different (13.1±2.78 vs. 16.1±3.11; *p*=0.47) (Fig. 4c-d). Final weights of mice in the early fate mapping groups were (27.8±1.4g vs. 49.2±0.8g; *p*=9.9e^−6^). To identify the cell fate of PDGFRα+ lineage, we labeled HA+ cells with oligodendrocyte lineage marker, Olig2. We found nearly 100% of the HA+ cells expressed Olig2, indicating that obesity does not alter the cell fate of PDGFRα+ OPC (Fig. 4e). These data indicate that the rate of OPC differentiation is accelerated during obesity development. In the absence of robust *in vivo* markers for post-differentiated but not completely mature myelinating oligodendrocytes, we turned to a gene expression approach to better characterize precisely when oligodendrocyte differentiation and maturation might be impaired by obesity.

**Figure 4.**
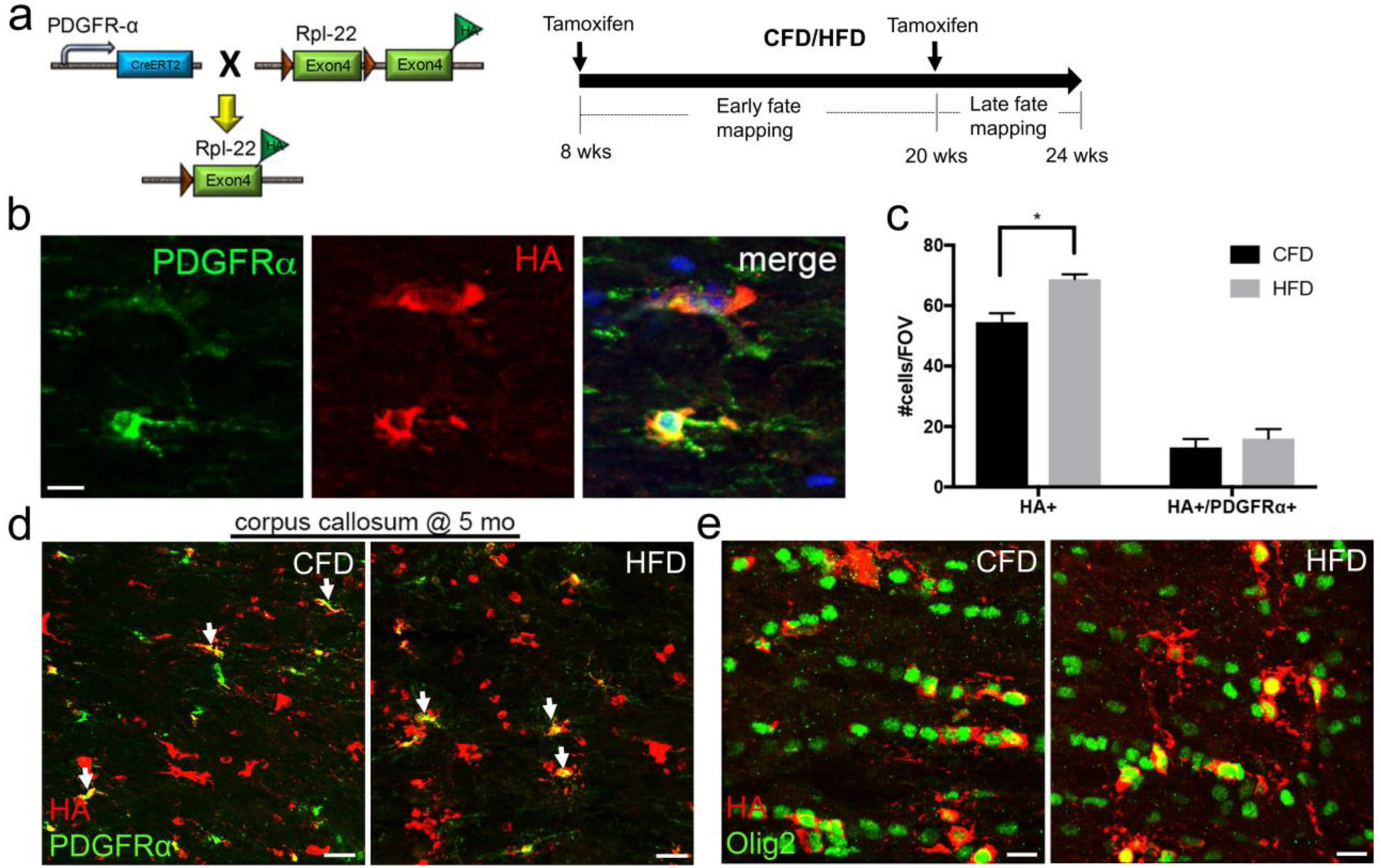
*PDGFRα-Cre^ERT^;Rpl22^tm1.1Psam^* fate mapping reveals accelerated OPC differentiation in obese mice. *PDGFRα*-*Cre^ERT^;Rpl22^tm1.1Psam^* mice result in tamoxifen-dependent Cre expression in OPCs and resultant Rpl22 exon 4 substitution with an HA-tagged exon 4 (a: left panel). Tamoxifen was administrated at two different time point for early and late fate mapping (a: right panel). After tamoxifen induction, there is robust HA labeling (red) within PDGFRα+ OPCs (green) that is largely restricted to the peri-nuclear region indicating ribosomal labeling (b; blue = dapi). Beginning at 8 weeks of age, Cre+:Rpl22-fl/fl mice (*n*=8) were fed a CFD or HFD for 12 weeks (*n*=4/grp). Tamoxifen induction was administered for 4d at the initiation of dietary change. Quantitation of HA+ and HA+/PDGFRα+ cells/fov (c). Co-labeling for HA and PDGFRα in the corpus callosum at 20 weeks shows animals on CFD have significant OPC differentiation over 12 weeks with 54.5% of cells/field of view HA+ and 13.1% of cells/fov HA+/PDGFRα+ (c-d). HA+ cells/fov were increased (68.8%) in animals on HFD with a similar number of HA+/PDGFRα+ cells/fov (16.1%) (c-d). Co-labeling for HA and Olig2 in the corpus callosum at 20 weeks shows nearly all HA+ cells express Olig2 in both CFD and HFD (e). Scale bar = 10 μm (b, e) and 20 μm (d). \**p*=0.011. Data are mean ± SEM.

### Obesity blocks myelinating oligodendrocyte gene expression

Together with our previous data of thinner myelin and shorter paranodal segments despite increased OPC differentiation, we reasoned that the maturation of oligodendrocyte was impeded by obesity. To determine at which stage the oligodendrocyte maturation block occurs, we developed a gene expression assay on the Nanostring^®^ platform using a probe set based on established markers of each stage of oligodendrocyte differentiation (OPC, pre-myelinating oligodendrocyte (PMO), and myelinating oligodendrocyte (MO) defined by cell type specific genes with FPKM > 20 (Zhang et al., 2014). The details for each probe set are available in the supplemental data file (Extended data Fig.5-1). All 120 genes were detectable using the assay and the expression of housekeeping genes was not significantly different between CFD and HFD. Differentially expressed genes (*p*<0.05) were classified as up- or down-regulated relative to mice on CFD. The number of differentially expressed genes per oligodendrocyte stage was determined and is shown in Figure 5. Among the significantly up-regulated genes (average *p*-value = 0.0226), 12/40 (30%) were OPC stage genes, 8/40 (20%) were PMO stage genes, and 1/40 (2.5%) were MO stage genes (Fig. 5a). Significantly more OPC genes were differentially expressed (Fisher Exact test; *p*=0.0108). The average log-fold change (log2FC) of the up-regulated genes were 0.35±0.17 (OPC), 0.23±0.09 (PMO) and 0.21 (MO) (Fig. 5b). Among the significantly down-regulated genes (average *p*-value = 0.017), all were MO stage genes 18/40 (45%) (Fig. 5c). The average log-fold change (log2FC) of the down-regulated MO genes were −0.26±0.10 (Fig. 5d). To evaluate the global gene expression profile of white matter from mice on CFD vs. HFD, we combined our complete gene expression data set with RPKM values for the marker genes of each oligodendrocyte stage from Zhang et al. (2014). Using tSNE, the similarity of CFD (red) and HFD (blue) white matter samples were compared to OPC, PMO, and MO (Fig. 5e). As expected, tSNE and hierarchical clustering (Fig. 5f) demonstrates that PMO (yellow) and MO (gray) gene expression profiles are quite similar, while OPCs (green) represent a distinct progenitor cell. CFD white matter samples cluster together as they represent a mix of all three oligodendrocyte cell types while HFD white matter samples cluster with the OPC population and away from the PMO and MO stages. Gene ontology analysis of the significantly different down-regulated genes indicates genetic programs in CNS and oligodendrocyte development, cytoskeletal organization, and myelination (arylsulfatase activity, GTP-Rho binding) are impacted by obesity. Gene ontology of up-regulated genes indicates genetic programs in anaerobic energy metabolism, production of extracellular matrix components, and signaling through PDGFRα are increased by obesity (Table 1). Gene ontology analysis using equal numbers of randomly selected genes from the probe set revealed no overlapping terms for either up- or down-regulated genes. Overall, these findings indicate that obesity impedes oligodendrocyte maturation at the transition between PMO and MO cell types accounting for the changes observed by light and electron microscopy.

**Figure 5.**
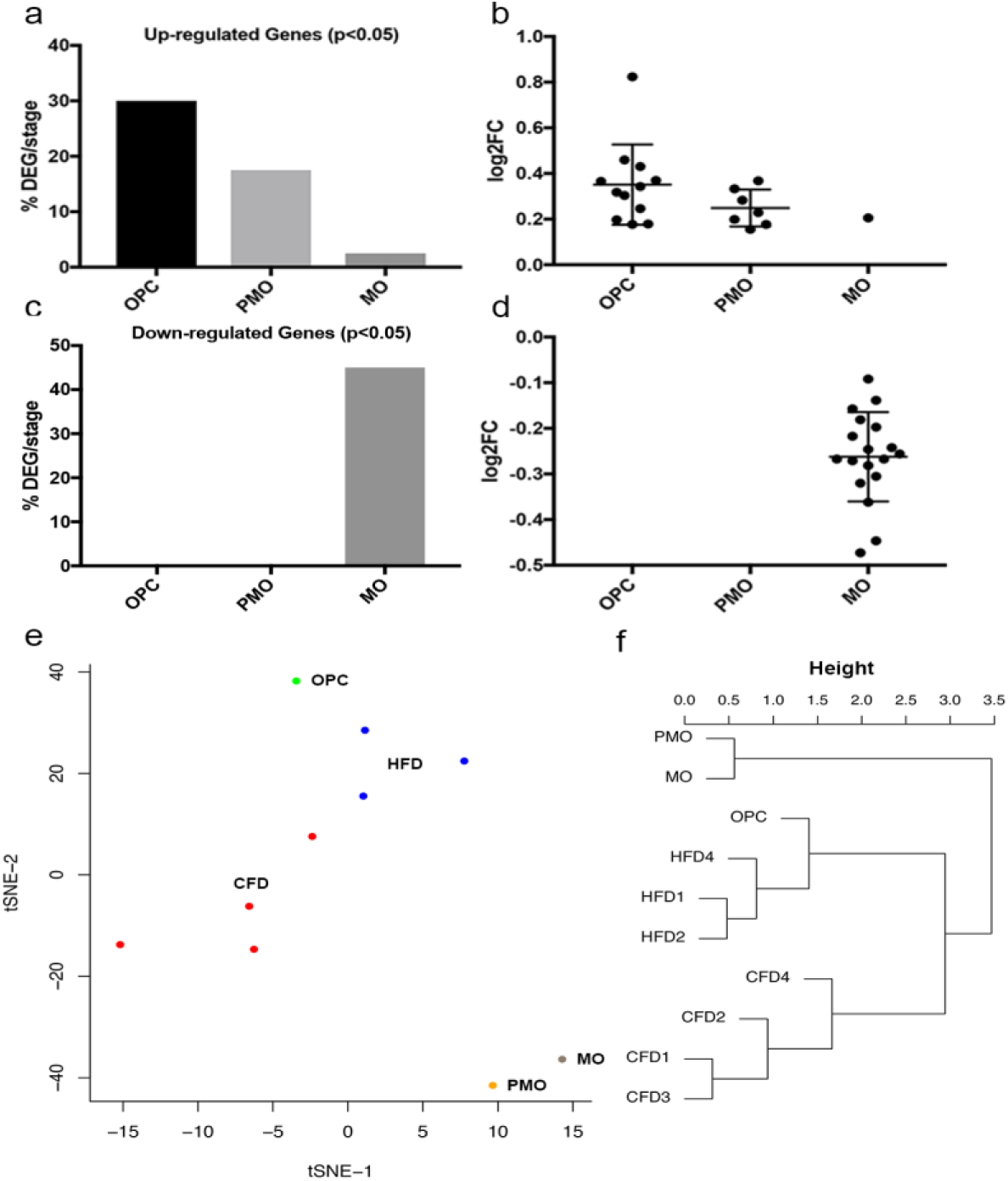
Oligodendrocyte staging by gene expression reveals obesity-induced maturation block. A Nanostring gene expression staging assay using known oligodendrocyte stage markers (40/stage) for OPC, premyelinating (PMO), and myelinating oligodendrocytes (MO) was used as described in Materials & Methods. Total RNA was isolated from callosal white matter from animals on CFD (*n*=4) or HFD (*n*=3) after 12 weeks on diet. Normalized counts of gene expression were compared for differential gene expression (*p*<0.05). Differentially expressed genes (DEGs) were enumerated by oligodendrocyte stage (%DEG/stage). Genes significantly up-regulated in animals on HFD were preferentially in the OPC (*p*=0.0018) and PMO stages (a). The average log-fold change (logFC) of the individual up-regulated DEGs are plotted by stage (b). Significantly down-regulated genes in animals on HFD were exclusively MO stage genes (c). The average log-fold change (logFC) of the individual down-regulated DEGs are plotted by stage (d). tSNE scatterplot of total gene expression profiles (120 genes) for animals on CFD (red) and HFD (blue) together with known OPC (green), PMO (yellow), and MO (gray) gene expression profiles (e). Hierarchical clustering of samples (f). logFC data are mean ± SEM.

**Table 1.**
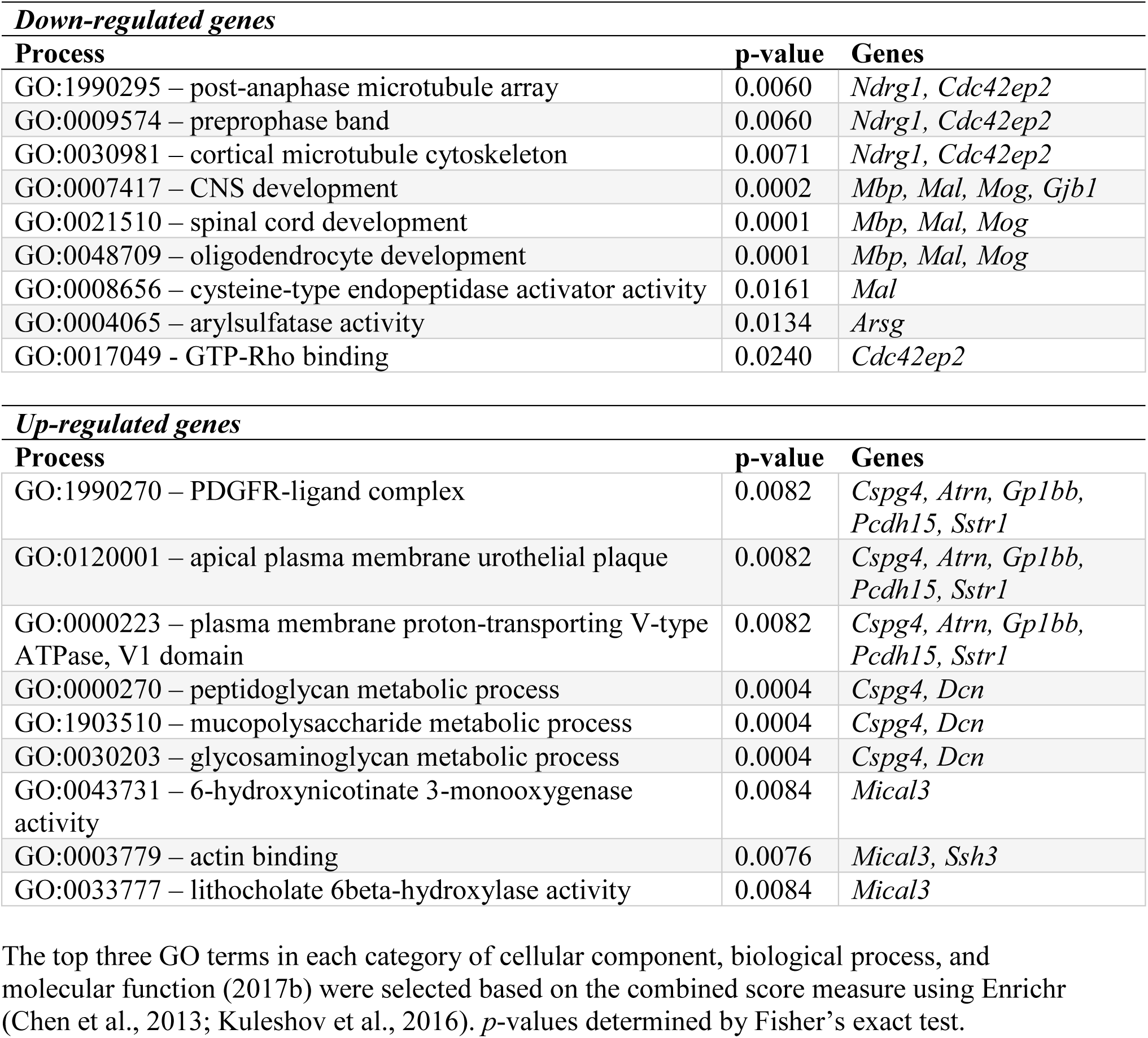
Gene ontologies for differentially expressed genes in HFD white matter

### Obesity-induced oligodendrocyte maturation block exacerbates early and impairs late responses to white matter stroke

Focal white matter lesions are characterized by dense loss of axons and oligodendrocytes, a robust inflammatory and astroglial response (Nunez et al. 2016), with stroke-responsive OPCs invading the lesion and playing a key role in repair and remyelination (Sozmen et al. 2016). Because we identified an obesity-induced oligodendrocyte maturation block, we reasoned that injury response and repair after white matter stroke were likely to be impaired by obesity. To test this hypothesis, we induced a focal white matter stroke and analyzed the acute response at 7d after stroke (Fig. 6) as well as the status of lesional repair at 28d after stroke (Fig. 7), when stroke-responsive OPCs are capable of limited differentiation, maturation and remyelination (Sozmen et al. 2016). At 7d after induction, focal white matter strokes can be identified by a dense loss of neurofilament and myelin basic protein staining (Fig. 6a). In animals on both CFD and HFD, labeling for surviving GST-π+ mature oligodendrocytes and stroke-responsive PDGFRα+ OPCs reveals a well-demarcated core lesion with little to no GST-π+ mature oligodendrocytes and a corresponding PDGFRα+ OPC rich area (Figs. 6b-c), though the average size of the stroke-responsive OPC area was larger in HFD animals (Fig. 6d; 180935±9459 μm^2^ vs. 235742±13371 μm^2^; *p*=0.002) consistent with an increased number of stroke-responsive OPCs (Fig. 6e; 137.3±21.17 vs. 247.7±14.5; *p*=0.013). The spatial localization of stroke-responsive OPCs were recorded in *x,y,z* coordinates relative to the center point of the stroke lesion (Fig. 6f). Nearest neighbor spatial analysis was used to compare stroke-responsive OPC distributions between animals on CFD and HFD (Fig. 6g). In lesion cores, the distribution of stroke-responsive OPCs were nearly identical between CFD (blue) and HFD (pink) (unshaded area). In contrast, at the lesion periphery, stroke-responsive OPCs were markedly more frequent in animals on HFD (pink shaded area) indicating an exaggerated early response to stroke in obese mice with the main effect on peri-infarct white matter.

**Figure 6.**
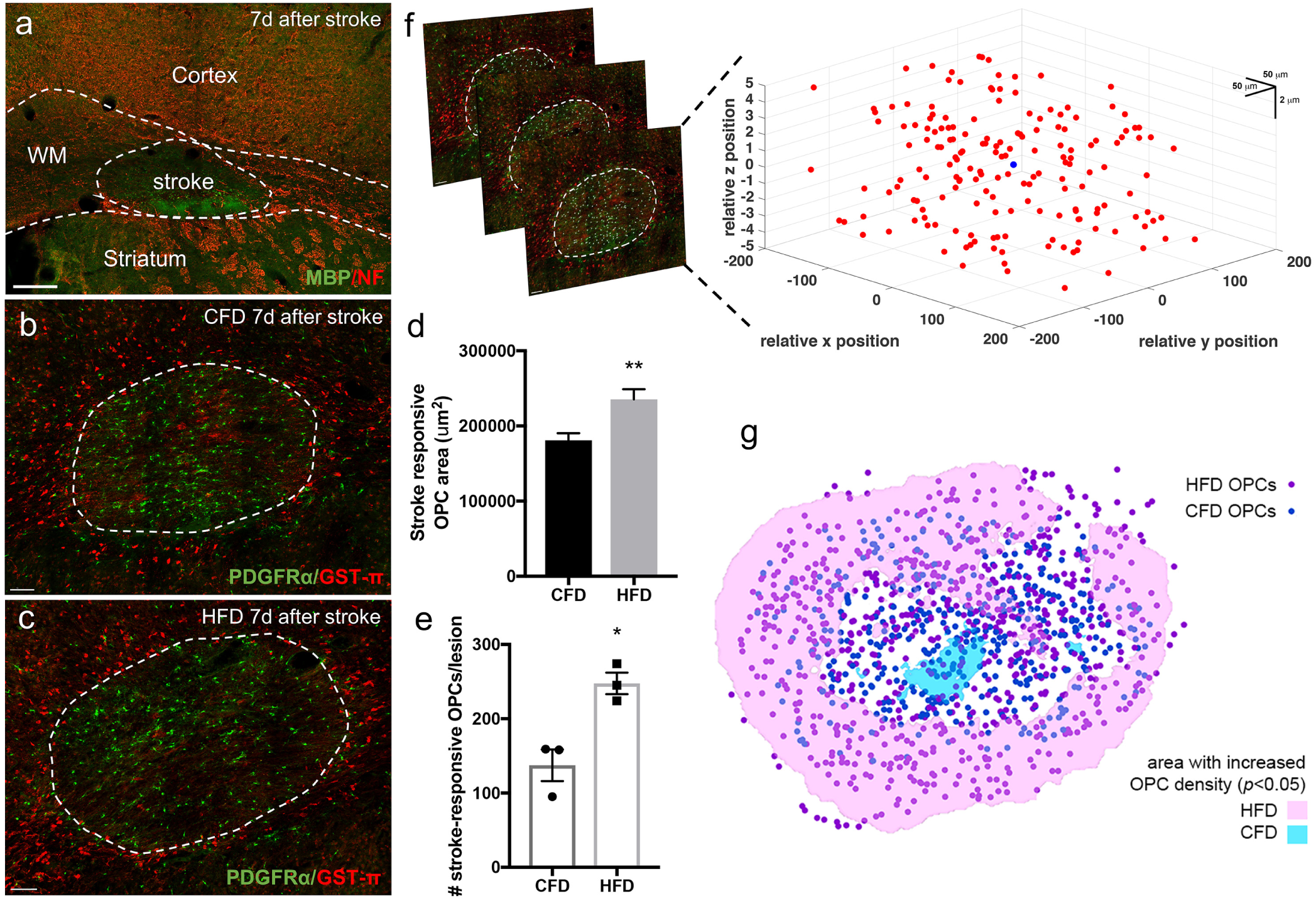
High fat diet induces an exaggerated OPC response to white matter stroke. White matter stroke induction in the left subcortical white matter underlying sensorimotor cortex produces a focal region of reduced myelin basic protein (MBP, green) staining and a loss of neurofilament-200 (NF, red) seven days after stroke (a). Labeling for GST-π+ mature oligodendrocytes (red) and PDGFR-α+ OPCs (green) in animals on CFD (*n*=4) (b) and HFD (*n*=4) (c) were used to identify stroke-responsive PDGFR-α+ OPCs (dashed line, b) and the extent of oligodendrocyte loss after stroke. Quantification of the stroke-responsive OPC area (µm^2^, d) and the number of stroke-responsive OPCs per lesion (e). Confocal z-stacks through the white matter stroke lesion were used and spatial coordinates (*x,y,z* in µm) relative to the center point of the stroke lesion (blue dot) were used to generate a spatial map of stroke-responsive OPCs (red) (f). Two-dimensional nearest neighbor spatial analysis comparing peri-infarct spatial frequency of stroke-responsive OPCs between CFD (blue) and HFD (pink) with the area of significantly (*p*<0.05) greater frequency of stroke-responsive OPCs between conditions in the shaded area (CFD, blue; HFD, pink) (g). Scale bar = 200 µm (a), 50 µm (d, e). Data are mean ± SEM.

To determine how this exaggerated early response in obesity impacts repair after stroke, we analyzed white matter stroke lesions 28d after stroke. At 28d after stroke, since PDGFRα+ OPCs underwent significant apoptosis, white matter stroke lesions are less well-demarcated by GST-π and PDGFRα labeling compared to the 7d time point (Fig. 7a). Therefore, we compared OPC and oligodendrocyte cell counts in three regions of interest across white matter in a central core lesion (middle box) and two lateral edge regions (left and right boxes) that correspond spatially with the area identified at 7d. This analysis revealed a significant change in oligodendrocyte cell populations 28d after stroke (*p*=0.0006, two-way ANOVA, *F*=9.854). Within the lesion core, a significantly higher number of PDGFRα+ OPCs were retained in animals on HFD compared to CFD (Fig. 7b-d; 9.67±0.84 vs. 17.45±1.40; *p*=0.034; *n*=3/grp). More OPCs were also detected in HFD animals at lesion edges (8.99±0.83 vs. 14.67±2.84; *p*=0.16). The morphologies of these cells were different from the typical bipolar OPCs morphology. Residual stroke-responsive OPCs in HFD animals adopted a more fibroblast morphology (Fig. 7c). Despite an increased frequency of PDGFRα+ OPCs at the lesion periphery at 7d after stroke, GST-π+ mature oligodendrocytes were equal at the edge of stroke (Fig. 7e-f, 7i; 15.00±1.42 vs. 15.50±2.50; *p*=0.99). Within the lesion core, a significantly lower number of GST-π+ mature oligodendrocytes were noted in animals on HFD (Fig. 7g-i; 17.98±1.25 vs. 7.44±2.39; *p*=0.004). These results indicate that the obesity-induced oligodendrocyte maturation block within white matter exaggerates the early response to stroke and subsequently limits the full maturation of oligodendrocytes after stroke.

**Figure 7.**
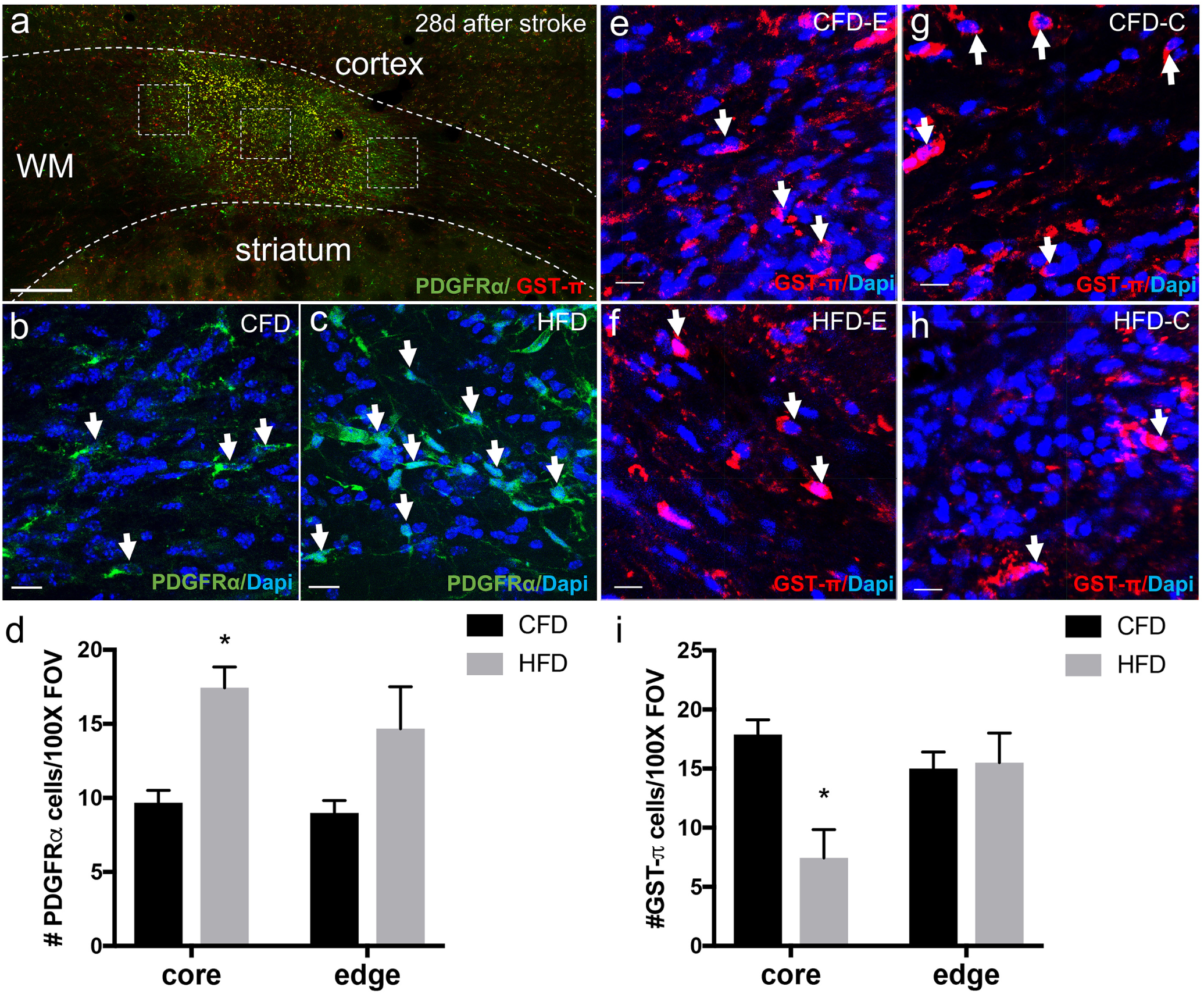
Obesity restricts the maturation of stroke-responsive OPCs. Twenty-eight days after white matter stroke induction labeling for GST-π+ mature oligodendrocytes (red) and PDGFR-α+ OPCs (green) within the stroke lesion identifies a lesion center and adjacent peri-infarct lesion edges (dashed inset boxes) (a). PDGFR-α+ OPCs within lesion center (CFD, b; HFD, c) are more numerous after HFD (d). GST-π+ mature oligodendrocytes within the peri-infarct edges (CFD, e; HFD, f) and lesion core (CFD, g; HFD, h) are reduced within the lesion center after HFD and equal at peri-infarct edges (i). Scale bar = 200 µm (a), 10 µm (b-c, e-h). \**p<*0.05. Data are mean ± SEM.

## Discussion

Obesity is linked to changes in brain white matter that rely on measures of structural integrity such as fractional anisotropy (Alfaro et al., 2018). The cellular consequences of obesity on white matter are not clear. In this study, after 12 weeks of diet-induced obesity, we found that cells of the oligodendrocyte lineage are specifically impaired. Adult onset obesity increases the resting number of OPCs within callosal white matter while accelerating OPC differentiation in an incomplete way. Despite increased rates of OPC differentiation, mice with adult onset diet-induced obesity have thinner myelin sheaths, increased g-ratios, shorter paranodal segments, and axonal microdomain clustering failures. Using gene expression analyses, adult onset obesity appears to impede oligodendrocyte maturation at or during the transition between pre-myelinating and myelinating oligodendrocyte stages, consistent with the ultrastructural changes observed by EM. In this obesity-primed state, white matter ischemic lesions produce an exaggerated OPC response leading to a measurable 30.1% increase in peri-infarct white matter in the early phase after stroke. During the remyelination and repair phase after white matter stroke (Sozmen et al., 2016), a persistent obesity-induced oligodendrocyte maturation block reduces mature oligodendrocytes within the core of the ischemic lesion. These findings indicate that adult onset diet-induced obesity exerts a specific effect on white matter cellular biology keeping OPCs in a progenitor state yet paradoxically impairs tissue recovery after stroke by blocking oligodendrocyte maturation and remyelination.

### The structural integrity of white matter is compromised by obesity

Observed reductions in fractional anisotropy seen in human imaging studies from subjects with obesity support a hypothesis of increased lateral water diffusivity through white matter tracts (Kullmann et al., 2016). The thinner myelin sheaths and resulting reduced axonal structural integrity in HFD mice can explain these findings. The approximate relationship between FA and the MR-estimated g-ratio is known and thought to be closely related approaching equivalence (Mohammadi et al., 2015). In the corpus callosum of obese subjects, FA declines by 9.6% (Karlsson et al., 2013), while here we observed an 8.75% increase in the g-ratio in callosal fibers, indicating that the approximate effect of HFD in mice on myelin relates in effect size to the change in FA seen in obese human subjects. Because we did not observe significant differences in fiber diameter, axonal diameter or axonal number by EM, the structural integrity of axons appears not be dramatically affected by obesity. However, declines in nodal and paranodal axonal complexes observed in HFD mice indicate that axonal conduction could be impaired. Single nucleotide polymorphisms in the neuronal cell adhesion gene, *Negr1*, are associated with obesity (Willer et al., 2009) and reduce FA in white matter tracts (Dennis et al., 2014) suggesting that there may also be neuronal factors driving reduced axoglial signaling and myelination in obese white matter.

### Potential mechanisms of obesity-induced oligodendrocyte block

Our results in adult onset diet-induced obesity are similar to those seen in genetically obese (*ob/ob*) mice with reductions in myelin (Sena et al., 1985) and increases in OPCs in leptin-deficient *ob/ob* mice (Udagawa et al., 2006). OPCs lack the leptin receptor after E18 and are not sensitive to leptin (Udagawa et al., 2006). Though increased in the peripheral circulation in HFD mice (Van Heek et al., 1997) and obese humans (Segal et al., 1996), leptin does not readily cross into the brain during obesity (Caro et al., 1996; Schwartz et al., 1996). Therefore, the effect of obesity on OPC number, differentiation and maturation is not due to leptin but more likely resulting from changes in metabolite availability or leptin-independent oligodendrocyte maturation pathways. Gene ontology results suggest that Rho family GTPase signaling is directly affected by obesity with Fyn and downstream Cdc42ep signaling a likely focal point for obesity-induced maturation block (Liang et al., 2004). Here, we used a marker gene expression profiling approach for OPC staging that, while able to clearly identify the stage of obesity-induced oligodendrocyte maturation block, fails to identify network signaling pathways that could be targeted to promote oligodendrocyte maturation during obesity. The increased numbers of PDGFRα-HA+ cells within callosal white matter during the development of adult onset obesity indicates that PDGFRα+ OPCs can efficiently exit to the PMO stage but cannot fully differentiate into mature myelinating oligodendrocytes. *MyRF* and *Fyn* kinase are key regulators of this transition (Umemori et al., 1994; Bujalka et al., 2013; Duncan et al., 2017) but the effect of obesity on these key regulatory checkpoints is not known. The lack of reliable markers for this PMO stage restricts the ability to enumerate these PMO cells. White matter primed with increased OPCs resulting from obesity is likely to impair signal transduction in white matter tracts vital for cognitive processing and can partially explain human imaging findings associated with obesity.

### Expansion of the white matter stroke penumbra in obesity

White matter ischemic lesions are associated with cognitive impairment and functional decline and increased by obesity (Alfaro et al., 2018). White matter ischemic lesions are characterized by a robust early loss of axons, myelin and oligodendrocytes (Valeriani et al., 2000; Back et al., 2007; Sozmen et al., 2009). Similar to inflammatory white matter lesions (Tripathi et al., 2010), OPCs respond early and robustly to white matter ischemic lesions (Sozmen et al., 2009; Miyamoto et al., 2015; Sozmen et al., 2016) though stroke-responsive OPCs largely adopt an astrocytic phenotype but can be stimulated to remyelinate by blocking Nogo receptor signaling (Sozmen et al., 2016). The peri-infarct white matter at the margin of the ischemic lesion, often referred to as the white matter penumbral region (Maillard et al., 2011), is where tissue repair is active. Axoglial signaling between axons and oligodendrocytes is compromised by brain ischemia (Reimer et al., 2011) and even more perilous in this white matter penumbral tissue (Hinman et al., 2015; Rosenzweig and Carmichael, 2015). Here, we used a novel approach to identify the spatial relationship of stroke-responsive OPCs to an elliptical lesion. This novel approach directly informs our data by showing that the increase in stroke-responsive OPCs produced by obesity occurs precisely in the peripheral margins of the white matter stroke lesion where tissue repair and axoglial signaling is maximal. Indeed, one month after stroke, obesity leaves behind a population of activated, injury-responsive OPCs whose maturation is inhibited. This progenitor restricted state induced by obesity could indicate that remyelination is simply delayed or it could lead to a progressive dysfunctional OPC response as the ability of NG2+ OPCs to differentiate into oligodendrocytes declines with chronic insults (Mason et al., 2004).

In conclusion, this study defines an obesity-induced oligodendrocyte maturation block state that alters the local response to and repair after focal white matter ischemia. Beyond ischemia, these findings have important bearing on the implementation of remyelination therapies. The efficacy of cellular and molecular remyelinating therapies deserves further study in the context of obesity if the translational potential of these treatment approaches is to be fully realized.

