## Supplementary Materials for "Obesity restricts oligodendrocyte maturation and impedes repair after white matter stroke" (from another version of this preprint: version 1, 2018-03-23)

| | CFD1 | CFD2 | CFD3 | CFD4 | HFD1 | HFD2 | HFD4 | Oligodendrocyte.Precursor.Cell | Newly.Formed.Oligodendrocyte | Myelinating.Oligodendrocytes |
|---|---|---|---|---|---|---|---|---|---|---|
| 1810041L15RIK | 58 | 58 | 55 | 60 | 61 | 58 | 59 | 65 | 95 | 60 |
| 9630013A20RIK | 9 | 7 | 7 | 6 | 7 | 6 | 6 | 22 | 100 | 75 |
| ACY3 | 7.5 | 8 | 8 | 9 | 9 | 8 | 9 | 2 | 10 | 63 |
| ADAP1 | 102 | 102 | 100 | 99 | 103 | 103 | 103 | 42 | 50 | 91.5 |
| ADSSL1 | 56 | 51 | 58 | 58 | 47 | 50 | 48 | 13 | 35 | 89 |
| APOD | 106 | 106 | 106 | 106 | 104 | 106 | 105 | 89 | 97 | 106 |
| ARSG | 55 | 49 | 64 | 52.5 | 43 | 46 | 44 | 23.5 | 30 | 76 |
| ASPA | 74 | 66.5 | 73 | 76 | 58 | 72 | 66 | 6 | 51 | 94 |
| ATRN | 96 | 94 | 94 | 95 | 98 | 98 | 99 | 46 | 56 | 57.5 |
| BMP4 | 6 | 13 | 9.5 | 7.5 | 8 | 10 | 8 | 43 | 58 | 33 |
| C1QL1 | 23 | 27 | 22.5 | 24 | 22 | 22 | 22 | 114 | 91 | 32 |
| CACNG4 | 24 | 26 | 26 | 27 | 25 | 23 | 27 | 112 | 86 | 30.5 |
| CARNS1 | 30 | 25 | 27 | 32 | 26 | 26 | 30 | 6 | 20 | 66 |
| CDC37L1 | 93 | 93 | 93 | 93 | 95 | 93 | 96 | 52 | 61 | 57.5 |
| CDC42EP2 | 37.5 | 35 | 34 | 36 | 27 | 34 | 29 | 10 | 38 | 86 |
| CDO1 | 50 | 52 | 53 | 50 | 54 | 53 | 49 | 108 | 47 | 27.5 |
| CDV3 | 89 | 89 | 89 | 91 | 92 | 90 | 92 | 54 | 80 | 53 |
| CENPB | 80 | 80 | 77 | 80 | 74 | 80 | 77 | 69 | 63 | 95 |
| CHN2 | 75 | 76 | 76 | 75 | 69 | 66 | 71 | 44 | 99 | 83 |
| CHRNA4 | 39 | 30 | 31 | 28 | 36 | 37 | 37 | 81 | 14 | 7.5 |
| CLDN11 | 101 | 101 | 102 | 102 | 100 | 101 | 101 | 39 | 114 | 114 |
| CNKSR3 | 11.5 | 12 | 16 | 13 | 11 | 11 | 12 | 32 | 64 | 61 |
| COL1A2 | 11.5 | 21 | 20 | 20 | 19 | 20 | 21 | 78 | 1.5 | 18 |
| CRYAB | 100 | 100 | 101 | 101 | 101 | 100 | 98 | 76 | 103 | 107 |
| CSPG4 | 22 | 23 | 25 | 22.5 | 24 | 25 | 28 | 103 | 43 | 7.5 |
| CYFIP2 | 114 | 114 | 114 | 114 | 114 | 114 | 114 | 79 | 89 | 49 |
| DCAF12L1 | 82 | 83 | 83 | 78 | 82 | 81 | 81 | 55.5 | 9 | 18 |
| DCN | 59 | 71 | 54 | 45 | 80 | 84 | 78 | 74.5 | 1.5 | 7.5 |
| DLL3 | 1 | 3 | 1 | 2 | 1 | 1 | 1 | 71.5 | 11 | 1.5 |
| EFHD1 | 60 | 57 | 59 | 61 | 48 | 54 | 47 | 17 | 45 | 90 |
| ELOVL6 | 44 | 46 | 45 | 44 | 46 | 44 | 46 | 51 | 65 | 62 |
| EMID1 | 7.5 | 6 | 6 | 5 | 6 | 7 | 7 | 104 | 48 | 22 |
| ENPP6 | 19 | 14 | 17 | 16 | 12 | 18 | 17 | 8 | 102 | 82 |
| FAM3C | 91 | 92 | 90 | 90 | 90 | 91 | 91 | 86 | 98 | 55 |
| FAM5C | 57 | 59 | 60 | 62 | 65 | 61 | 62 | 92 | 37 | 23.5 |
| FAM70B | 4 | 5 | 4 | 4 | 5 | 5 | 5 | 101 | 40 | 18 |
| FAM73A | 87 | 88 | 88 | 87 | 86 | 89 | 88 | 40.5 | 49 | 43 |
| FMNL2 | 95 | 95 | 95 | 96 | 97 | 96 | 97 | 53 | 62 | 56 |
| FRMD4A | 34 | 38 | 35 | 29 | 39 | 41 | 42 | 27 | 76 | 44 |
| FYN | 103 | 103 | 103 | 103 | 105 | 104 | 104 | 97 | 109 | 87 |
| GJB1 | 32 | 31 | 30 | 33 | 23 | 30 | 26 | 40.5 | 55 | 97 |
| GLRB | 109 | 109 | 109 | 109 | 108 | 108 | 108 | 80 | 88 | 70 |
| GP1BB | 14.5 | 10 | 11 | 11 | 15 | 12 | 13 | 38 | 84 | 45 |
| GRIA3 | 108 | 108 | 108 | 108 | 109 | 109 | 109 | 88 | 32 | 18 |
| GRIN3A | 63 | 79 | 66 | 72 | 72 | 71 | 74 | 62.5 | 8 | 3 |
| GSN | 92 | 91 | 96 | 92 | 91 | 92 | 89 | 95 | 108 | 109 |
| HAPLN2 | 28 | 29 | 28 | 37 | 30 | 29 | 33 | 2 | 27 | 80 |
| HCN2 | 104 | 104 | 104 | 105 | 102 | 102 | 102 | 48 | 41 | 88 |
| INF2 | 48 | 48 | 48 | 51 | 49 | 48 | 53 | 25 | 33 | 81 |
| INSC | 5 | 4 | 5 | 7.5 | 4 | 4 | 4 | 14 | 23 | 68 |
| ITGB4 | 21 | 20 | 21 | 21 | 18 | 21 | 20 | 2 | 16 | 71 |
| ITPR2 | 35.5 | 37 | 39 | 42 | 42 | 38 | 45 | 49 | 59 | 38 |
| KCNK1 | 33 | 32 | 36 | 26 | 35 | 32 | 32 | 64 | 6 | 25 |
| KIF19A | 10 | 9 | 9.5 | 10 | 10 | 9 | 10 | 18 | 68 | 50 |
| KIF5A | 115 | 115 | 115 | 115 | 115 | 115 | 115 | 66 | 70 | 99 |
| LHFPL3 | 52 | 55 | 56 | 59 | 53 | 49 | 51 | 110 | 77 | 34 |
| LIMS2 | 40 | 39.5 | 41 | 39 | 37 | 43 | 38 | 99 | 104 | 84 |
| LNX1 | 13 | 16 | 12 | 14 | 16 | 17 | 16 | 93 | 19 | 13 |
| LYPD6 | 35.5 | 33 | 29 | 30 | 34 | 33 | 34 | 61 | 7 | 27.5 |
| MAL | 107 | 107 | 107 | 107 | 107 | 107 | 107 | 26 | 113 | 113 |
| MBP | 111 | 111 | 110 | 112 | 111 | 111 | 111 | 111 | 115 | 115 |
| MEGF11 | 31 | 34 | 33 | 35 | 40 | 35 | 43 | 84 | 28.5 | 7.5 |
| MICAL3 | 62 | 64 | 63 | 66 | 71 | 69 | 79 | 23.5 | 52 | 42 |
| MMP15 | 26 | 24 | 37 | 31 | 29 | 24 | 23 | 109 | 57 | 41 |
| MOG | 98 | 96 | 99 | 100 | 93 | 97 | 94 | 21 | 110 | 110 |
| MYCL1 | 66 | 68 | 62 | 65 | 63 | 64 | 61 | 59 | 74 | 39 |
| MYT1 | 42 | 45 | 42 | 49 | 51 | 47 | 56 | 82 | 22 | 7.5 |
| NCALD | 113 | 112 | 112 | 111 | 113 | 113 | 113 | 107 | 67 | 46 |
| NDRG1 | 97 | 97 | 97 | 97 | 94 | 94 | 95 | 11.5 | 66 | 103 |
| NFASC | 77 | 77 | 78 | 81 | 78 | 83 | 84 | 50 | 111 | 96 |

| | | | | | | | | | | |
|---|---|---|---|---|---|---|---|---|---|---|
| NOL3 | 18 | 19 | 19 | 19 | 21 | 19 | 19 | 20 | 15 | 64 |
| NXPH1 | 83 | 85 | 80 | 84 | 87 | 86 | 85 | 102 | 42 | 13 |
| OPALIN | 81 | 75 | 79 | 82 | 56 | 73 | 60 | 6 | 90 | 105 |
| PCDH15 | 45 | 42 | 44 | 40 | 50 | 52 | 54 | 83 | 17 | 18 |
| PDGFRA | 71 | 61 | 69 | 63 | 66 | 62 | 63 | 115 | 87 | 23.5 |
| PDLIM2 | 42 | 39.5 | 43 | 46 | 33 | 36 | 36 | 16 | 75 | 101 |
| PID1 | 53 | 50 | 52 | 56 | 55 | 56 | 55 | 94 | 31 | 13 |
| PIK3R3 | 64.5 | 62 | 65 | 64 | 67 | 63 | 65 | 36.5 | 85 | 48 |
| PLA2G16 | 46 | 44 | 47 | 41 | 41 | 42 | 39 | 15 | 18 | 74 |
| PLEKHB1 | 112 | 113 | 113 | 113 | 112 | 112 | 112 | 105 | 112 | 112 |
| PNLIP | 3 | 1 | 2.5 | 1 | 3 | 3 | 2 | 58 | 5 | 1.5 |
| PPAPDC1A | 25 | 22 | 22.5 | 22.5 | 28 | 28 | 25 | 100 | 39 | 18 |
| PPP1R14A | 47 | 47 | 49 | 52.5 | 44 | 45 | 40 | 9 | 71 | 104 |
| PPP2R3A | 69.5 | 69 | 72 | 68 | 64 | 67 | 70 | 35 | 44 | 47 |
| PRR18 | 84 | 81 | 82 | 83 | 75 | 82 | 75 | 34 | 72 | 100 |
| RAP2A | 105 | 105 | 105 | 104 | 106 | 105 | 106 | 77 | 101 | 85 |
| RASGRF1 | 67 | 72 | 85 | 70 | 85 | 68 | 69 | 60 | 3.5 | 27.5 |
| RFTN1 | 29 | 36 | 32 | 34 | 31 | 31 | 35 | 11.5 | 12 | 69 |
| RLBP1 | 42 | 41 | 38 | 38 | 38 | 39 | 31 | 106 | 46 | 27.5 |
| RNF122 | 14.5 | 15 | 15 | 17 | 14 | 13 | 14 | 47 | 82 | 73 |
| RPRM | 68 | 66.5 | 57 | 57 | 77 | 74 | 67 | 91 | 26 | 36 |
| RRAS2 | 64.5 | 65 | 68 | 67 | 70 | 70 | 73 | 33 | 92 | 78 |
| SAMD4B | 51 | 56 | 50 | 48 | 52 | 51 | 50 | 67 | 73 | 65 |
| SDC3 | 69.5 | 74 | 67 | 71 | 73 | 75 | 72 | 113 | 83 | 37 |
| SEMA5A | 73 | 70 | 71 | 73 | 68 | 65 | 68 | 71.5 | 79 | 59 |
| SHC4 | 17 | 18 | 13 | 15 | 13 | 14 | 15 | 73 | 13 | 18 |
| SLC12A2 | 76 | 63 | 81 | 74 | 60 | 60 | 58 | 57 | 93.5 | 79 |
| SLC45A3 | 2 | 2 | 2.5 | 3 | 2 | 2 | 3 | 4 | 21 | 67 |
| SLCO3A1 | 72 | 73 | 70 | 69 | 76 | 76 | 76 | 30 | 25 | 77 |
| SLITRK1 | 88 | 86 | 86 | 85 | 89 | 88 | 90 | 62.5 | 3.5 | 7.5 |
| SMOC1 | 27 | 28 | 24 | 25 | 32 | 27 | 24 | 87 | 28.5 | 7.5 |
| SSH3 | 54 | 53 | 51 | 54 | 57 | 57 | 64 | 19 | 69 | 52 |
| SSTR1 | 49 | 54 | 46 | 43 | 59 | 55 | 52 | 90 | 24 | 7.5 |
| STRN | 78 | 78 | 74 | 79 | 79 | 77 | 80 | 29 | 53 | 51 |
| TBC1D9B | 99 | 99 | 98 | 98 | 99 | 99 | 100 | 55.5 | 54 | 93 |
| TMEM108 | 20 | 17 | 18 | 18 | 20 | 16 | 18 | 28 | 60 | 40 |
| TMEM163 | 37.5 | 43 | 40 | 47 | 45 | 40 | 41 | 85 | 106 | 91.5 |
| TMEM179 | 79 | 82 | 75 | 77 | 81 | 78 | 82 | 98 | 34 | 35 |
| TNS3 | 86 | 87 | 84 | 86 | 83 | 85 | 86 | 74.5 | 96 | 72 |
| TPPP3 | 85 | 84 | 87 | 88 | 84 | 79 | 83 | 31 | 78 | 102 |
| TRAK2 | 94 | 98 | 92 | 94 | 96 | 95 | 93 | 36.5 | 81 | 98 |
| TRF | 110 | 110 | 111 | 110 | 110 | 110 | 110 | 70 | 107 | 111 |
| TRP53INP2 | 90 | 90 | 91 | 89 | 88 | 87 | 87 | 68 | 105 | 108 |
| UGDH | 16 | 11 | 14 | 12 | 17 | 15 | 11 | 96 | 36 | 30.5 |
| UST | 61 | 60 | 61 | 55 | 62 | 59 | 57 | 45 | 93.5 | 54 |
