## Supplementary Materials for "Obesity restricts oligodendrocyte maturation and impedes repair after white matter stroke" (from another version of this preprint: version 1, 2018-03-23)

| | Customer Ide | Accession | Position | Target Sequence | Tm CP | Tm RP | Flags | Stage | HUGO Gene | NSID | Design Remarks |
|---|---|---|---|---|---|---|---|---|---|---|---|
| 1 | Pdgfra | NM_001083316.1 | 2521-2620 | CTCAGAAGTCAAAAACCTCCTTTCGGACGATGACTCCGAGGGTCTGACTTTGCTGGATCTATTGAGCTTCACCTATCAAGTTGCTCGAGGAATGGAATTT | 82 | 79 | | OPC | Pdgfra | NM_001083316.1:2520 | |
| 2 | Lnx1 | NM_010727.4 | 1251-1350 | GCTGGAGGAGAATGACCGTGTGCTGGCCATCAACGGGCATGACCTTCGATTTGGCAGTCCAGAGAGTGCAGCTCATCTGATTCAGGCCAGTGAAAGACGC | 82 | 81 | | OPC | Lnx1 | NM_010727.4:1250 | targets 2nd PDZ domain conserved in all 5 transcripts |
| 3 | Dcn | NM_001190451.2 | 1001-1100 | CAAGGTTGATGCACCCAGCCTGAAAGGACTGATTAATTTGTCTAAACTGGGATTGAGCTTCAACAGCATCACCGTTATGGAGAATGGCAGTCTGGCCAAT | 82 | 82 | | OPC | Dcn | NM_001190451.2:1000 | targets exon6-7 junction in CDS |
| 4 | Mmp15 | NM_008609.3 | 1766-1865 | CCGGGGAACATCAGTGCTGCCTACGAGCGCCAGGATGGACATTTTGTCTTCTTCAAAGGTAACCGCTACTGGCTTTTCCGAGAAGCCAATCTGGAGCCCG | 82 | 82 | | OPC | Mmp15 | NM_008609.3:1765 | |
| 5 | Cdo1 | NM_033037.3 | 693-792 | CACGCCTTTGACCAGAGAACAGGGCATAAAAACAAAGTCACCATGACATTCCACAGCAAGTTTGGAATCAGGACTCCATTTACAACTTCAGGGTCACTGG | 82 | 82 | | OPC | Cdo1 | NM_033037.3:692 | |
| 6 | Sapcd2 | NM_001081085.1 | 1519-1618 | AGACTTGGGTGCTGCTAGAAATGTACTCTGCCGGAAGCTGCCACTTGCCACTGTCAGTGGGTGGCGGAGGATCATCCTCACGTTTACCTGATTTCAGGCA | 83 | 83 | | OPC | Sapcd2 | NM_001081085.1:1518 | |
| 7 | Kcnk1 | NM_008430.2 | 957-1056 | CCGCCGTGTTCTCTGTGCTGGAGGATGACTGGAACTTCCTCGAGTCTTTTTACTTCTGTTTCATCTCCCTGAGCACCATCGGCCTGGGGGACTATGTTCC | 82 | 82 | | OPC | Kcnk1 | NM_008430.2:956 | |
| 8 | Rasgrf1 | NM_011245.2 | 159-258 | CGAGGGGCATTCGGATGCCTCTCTGAGATGTAAGTGCGTAGCAGTCTCTCAGGCCTTCTAGCATCATGCAGAAAGCCATCCGGCTTAACGATGGCCACGT | 82 | 81 | | OPC | Rasgrf1 | NM_011245.2:158 | |
| 9 | Pcdh15 | NM_001142741.1 | 623-722 | GTTAAACAGATGCTTTTCCTGAACAGTACCGGAAGAGTTCTGGATAGAGACCCACCAATGAACATACACTCCATTGTGGTGCAAGTCCAGTGTGTCAACA | 80 | 82 | | OPC | Pcdh15 | NM_001142741.1:622 | |
| 10 | Chrna4 | NM_015730.5 | 2876-2975 | CCTTCAATGTTGTAGAATAGCGTGAGCTGCCAAGAGACTTCTAAGCAAAACAGGCTCTGTGACTCATTTTTCGAGGGCCATCGACCAAGTCTTAGGGGTG | 80 | 82 | | OPC | Chrna4 | NM_015730.5:2875 | |
| 11 | Dll3 | NM_007866.2 | 840-939 | ATGCCTTTTACCTGGGCCTGGACCTTGTGATGGGAACCCATGTGCCAATGGGGGCAGCTGTAGTGAAACCTCTGGCTCCTTTGAATGTGCCTGTCCCCGG | 85 | 84 | | OPC | Dll3 | NM_007866.2:839 | |
| 12 | Col1a2 | NM_007743.2 | 4026-4125 | GCTGTGCTTCTGCAGGGTTCCAACGATGTTGAACTTGTTGCTGAGGGCAACAGCAGGTTCACCTACTCTGTCCTAGTCGATGGCTGCTCCAAAAAGACAA | 82 | 83 | | OPC | Col1a2 | NM_007743.2:4025 | |
| 13 | Fam70b | NM_001143671.1 | 741-840 | ACCGGCTGCTCTGGGTATCCACAGTGCTGAACGTATTGGGCCTGTGCCTGGGCATTGTCACAGCTGCAGTCCTAGGGGCCTTCAAGGACATGGTCCCTCT | 82 | 83 | | OPC | Tmem255b | NM_001143671.1:740 | |
| 14 | Sstr1 | NM_009216.3 | 2046-2145 | CAAGACAGTCCCTAATCACCGCTCTCCTTTCGCAGAGCCTTACTGTCAAGGAAGCCCAGCTCGAGGATGTCTAGGCAACTTGTCTTTTCTACTCTCAGAG | 82 | 83 | | OPC | Sstr1 | NM_009216.3:2045 | |
| 15 | Pnlip | NM_026925.3 | 883-982 | GTTTTACACCGACAGCATCGTCAACCCAACTGGCTTTGCTGGGTTCTCCTGTAGCTCTTACAGTCTTTTCACGGCAAATAAGTGCTTCCCATGTGGGAGT | 81 | 79 | | OPC | Pnlip | NM_026925.3:882 | |
| 16 | Cspg4 | NM_139001.2 | 1531-1630 | TGGAGCTAGACATCCTGGGAGCCCAAACCCGGAAAATGTTTACCCTGTTGGACGTGGTGAACCGTAAGGCCCGCTTTGTTCACGATGGCTCTGAAGACAC | 83 | 83 | | OPC | Cspg4 | NM_139001.2:1530 | |
| 17 | Lppr1 | NM_178756.4 | 971-1070 | TGATCGCCGGCTTCATCCTGGGCACCGCAGTGGCCCTGTTTTTGGGAATGTGTGTGGTTCATAACTTCAGAGGAACCCAAGGCTCTCCTTCCAAACCCAA | 82 | 82 | | OPC | Plppr1 | NM_178756.4:970 | |
| 18 | Ppapdc1a | NM_001080963.1 | 673-772 | CCTATATTTGCTACAGACAACATTATCCTCCTCTGGCCAACACAGCATGCCACAAACCGTATGTCAGCCTTCGAGTGCCCACTTCACTGAAGAAGGAAGA | 83 | 82 | | OPC | Plpp4 | NM_001080963.1:672 | |
| 19 | Nxph1 | NM_008751.5 | 1161-1260 | GCTCTTGTTGACTGGTGCACACATGCTAGTGGAAAACAATTGATGTCATTTCTGCACAGTCAGCTTCATCTCTCAGCATAGTTGTAAATCACCATAGATC | 80 | 78 | | OPC | Nxph1 | NM_008751.5:1160 | |
| 20 | Pid1 | NM_001003948.2 | 420-519 | CTGATGAAGGCAAGGACTCATAGTGGCTGCAAGGTTACCTATCTGGGAAAAGTCTCTACCACAGGCATGCAGTTTTTGTCTGGTTGCACCGAGAAGCCAG | 83 | 82 | | OPC | Pid1 | NM_001003948.2:419 | |
| 21 | Ugdh | NM_009466.2 | 731-830 | ATCAAGGACCTAAAGAACCCGGACAGAGTCCTGATTGGAGGGGATGAAACTCCAGAGGGCCAGAAAGCCGTTCGGGCTCTGTGTGCTGTGTATGAGCACT | 82 | 82 | | OPC | Ugdh | NM_009466.2:730 | |
| 22 | Slitrk1 | NM_199065.2 | 851-950 | TTCTCTGGGATCGGACATTGGAGCACTAAATGAACTTGAATTGTGTCTGTGGCGAGCAGGATGGTTGCTGTTACTTTGTGATGAGATCTGGGATGAATTG | 80 | 82 | | OPC | Slitrk1 | NM_199065.2:850 | |
| 23 | Shc4 | NM_199022.2 | 2945-3044 | TTTCAGATGCTAAAGACAACTTGAACTGCTCAGGCAGTGGTACGTGCCTTTACCTTGATAATCCGGGGACATTTTCCTATGGGGGAGATTAGTTACACGT | 82 | 79 | | OPC | Shc4 | NM_199022.2:2944 | |
| 24 | Smoc1 | NM_001146217.1 | 2956-3055 | TGGTTCTTTTGCTGGCTTGCTCTCCTCAACCTGCCTTCCCTGGGTTTCCATCTCAGTATCCTGTAAATGCATCCACCCTGGTAAACCCTGCTCTATTTTG | 81 | 79 | | OPC | Smoc1 | NM_001146217.1:2955 | |
| 25 | Emid1 | NM_080595.2 | 1129-1228 | GGGTTGCACCAACTACGCGAGGCTTTGAAGATTTTAGCTGAGAGGGTTTTAATCTTGGAAACAATGATTGGGCTCTATGAACCAGACTTGGGATCCGGGG | 80 | 82 | | OPC | Emid1 | NM_080595.2:1128 | |
| 26 | Rlbp1 | NM_001173483.1 | 1453-1552 | CCGTATTAGCATCCCAGTCCTTCTTCAGCTTTCCTGAACCCAAGTCCGGGGACAGGAATGCCTGAAGGGACTGTTTTCCCAGACTTCCCTAGGTTTAGGA | 80 | 82 | | OPC | Rlbp1 | NM_001173483.1:1452 | |
| 27 | Dcaf12l1 | NM_001190718.1 | 2911-3010 | GAGCTGGGGATTAAAAAGATAGGTCTGATTAAAGGACTGTGCAGTTCAGATAAATAGAGATAATGGGATGCCGTGCGGATAGATTGGTTTTCTCCCTAGG | 82 | 80 | | OPC | Dcaf12l1 | NM_001190718.1:2910 | |
| 28 | Lypd6 | NM_177139.5 | 1551-1650 | CAGGTGTTGCATTCTAGTCAGATATAGCTGATGTAGTCAGAGAGAGGACTTCAGAATAAAAGGAAGCTTTCTGGAGGTTAGTCCTAGCTCATTTGGAAGG | 82 | 80 | | OPC | Lypd6 | NM_177139.5:1550 | |
| 29 | Lhfpl3 | NM_029990.1 | 2341-2440 | CCCAAAGCACTAAATTAAGGGAATGTTAGCCTCTGTAATTGTAGTCCTGGTCATCCTGGGGGCTGGTCATTGCTTTAGGTACTTGTCCCGAGGCATTCAG | 79 | 82 | | OPC | Lhfpl3 | NM_029990.1:2340 | |
| 30 | Myt1 | NM_008665.4 | 1441-1540 | TCAGTCATCGTGGAGGTTCGCTCTGATGATGACAAGGATGAGGATTCACGCTCCCAGAAGTCAGCAGTCACAGATGAATCAGAAATGTATGACATGATGA | 83 | 83 | | OPC | Myt1 | NM_008665.4:1440 | |
| 31 | Gria3 | NM_016886.3 | 391-490 | GCTTTCCGCTTTGCTGTGCAGTTATACAACACCAACCAGAACACCACTGAGAAGCCCTTCCATTTGAACTACCACGTAGACCACTTGGATTCCTCCAATA | 83 | 81 | | OPC | Gria3 | NM_016886.3:390 | |
| 32 | C1ql1 | NM_011795.2 | 963-1062 | CCCTCTTCCCGCACGCACGCAGTTCCGGCCCGGGTGGCAGGTTGCCTTCCTGGTCCCAGCCTGCACTGTATATGTGTACAATAGGACTGTTTACTGAGCC | 85 | 85 | | OPC | C1ql1 | NM_011795.2:962 | |
| 33 | Tmem179 | NM_178915.3 | 389-488 | CTCTTCTTCCTCTGCAAGGGACACGAGGGCTCCTTCTTCTACGCCTTCCTGAACCTCCTGGTCAGTGCCTTTGTGGTCTTCCTGGTCTTCATTGCCAGCA | 82 | 82 | | OPC | Tmem179 | NM_178915.3:388 | |
| 34 | Megf11 | NM_001134399.1 | 345-444 | GATCAGATCTACTACACACGATGTGCAGACATCCTCAACTGGTTCAAGTGTACCCGGCACCGGATCAGCTATAAGACCGCGTATAGGCGCGGCCTCCGGA | 83 | 82 | | OPC | Megf11 | NM_001134399.1:344 | |
| 35 | Ncald | NM_134094.3 | 1416-1515 | GTTTGAAATACAAAACTCCTACCCTGGGTCTGTTGTCACACATGCCCTGGGAGCTAGAGCAGACACCTGAGGCTAAGTTTTTAAAGAACAGGTTACCCTC | 83 | 82 | | OPC | Ncald | NM_134094.3:1415 | |
| 36 | Sdc3 | NM_011520.3 | 2276-2375 | CACTGTCATTGTCACCAGTGGCGACCACACCACCTTCCTCCCAATCTATTCTCTTTACACACATTGGTCCTGCATCTGGCTCCTCTAACCATCCAGGTCA | 83 | 82 | | OPC | Sdc3 | NM_011520.3:2275 | |
| 37 | Rprm | NM_023396.5 | 995-1094 | CTGTTTATTCGCCACGCTGCAGACCTAATTGGGAAGGAACATAGATGCGTGTGGGTTTTGCGACCTACTAAGAGAGTGAAACTTTGATCGATAGTAACCT | 82 | 79 | | OPC | Rprm | NM_023396.5:994 | |
| 38 | Cacng4 | NM_019431.2 | 525-624 | AAAGAACCATTACAACTACGGCTGGTCTTTTTACTTTGGAGCCCTGTCGTTTATTGTGGCGGAGACCGTGGGCGTCCTGGCTGTAAACATTTACATTGAG | 78 | 82 | | OPC | Cacng4 | NM_019431.2:524 | |
| 39 | Grin3a | NM_001033351.1 | 1333-1432 | CTCAGTGGTTCCATCAAAGTAAAGGGATCCACCATCGTCAGCTCAGAAAACAACTTTTTCATCTGGAACTTGCAGTATGACCCTATGGGAAAGCCAATGT | 82 | 79 | | OPC | Grin3a | NM_001033351.1:1332 | |
| 40 | Fam5c | NM_153539.3 | 1156-1255 | CAGAGTCCTGAGAATAAAATTCAGTTGCAAGGGCTTCAAGTTCTCCTTCCCGACTACCTTCAGGAACGTTTTGTACAGGCAGCACTGAGCTACATTGCTT | 78 | 81 | | OPC | Brinp3 | NM_153539.3:1155 | |
| 41 | Gp1bb | NM_010327.2 | 1556-1655 | ACCCGAGCTCTGCAAGCCTGAATCAACATGGTGCCAAACTGATTCCAGTACTACCAGAGTGAAGCAACCTTGAGTCTTCAGAGCCTAGAGTGATGGAACA | 82 | 83 | | PMO | Gp1bb | NM_010327.2:1555 | |
| 42 | Tmem108 | NM_178638.4 | 2285-2384 | AAGCTGAACGTTAGTGTGGCCAAGTGAGTGGTCATCTGCGTGGTCTTGATGCTCTGGTTCCCAGGACTCAGTGTGCCCTCATCTCCTGTCTTTTTGTATG | 81 | 79 | | PMO | Tmem108 | NM_178638.4:2284 | |
| 43 | Fyn | NM_008054.2 | 1031-1130 | TTCTTATCCGCGAGAGCGAAACCACCAAAGGTGCCTACTCACTTTCCATCCGTGATTGGGATGATATGAAAGGGGACCACGTCAAACATTATAAAATCCG | 82 | 80 | | PMO | Fyn | NM_008054.2:1030 | |
| 44 | Ust | NM_177387.3 | 2445-2544 | TGATATGTGTGGCAGGAATCCTTAGCTCAATTGCACACCGTGTGCGATCCTGAAAGCACACTCATAGCCAATGGGGCTATACTGAACTTGGCATTTTTCA | 81 | 81 | | PMO | Ust | NM_177387.3:2444 | |
| 45 | Mical3 | NM_001270475.1 | 627-726 | GCACAGCCATCGACTTGTCCTTGTTGGGAGCCAAGGTGGTTGTTATTGAAAAGCGAGATGCCTTCTCCCGCAACAATGTCTTACATCTCTGGCCCTTCAC | 81 | 82 | | PMO | Mical3 | NM_001270475.1:626 | |
| 46 | Kif19a | NM_001102615.1 | 2903-3002 | TGGGCGGCCATCTCCTGGTGTCAGGCATCTGGGAAAAGTCTCTCTGCCTCTGGCCAAGGTCAAATTTCCTCCAAACCAGAACACAGGATCTGGAAACCCT | 83 | 82 | | PMO | Kif19a | NM_001102615.1:2902 | |
| 47 | 1810041L15R | NM_001163145.1 | 3857-3956 | TGGCTTTTTCCCAGTGGGAACTCTATGTCCCTCGTGATCAGTGTTACGCCCTGTCATCTGTGAGCTCTGGAACCTCCATTCAGAACTCCTCCCTGGTTGG | 81 | 81 | | PMO | 1810041L15 | NM_001163145.1:3856 | |
| 48 | 9630013A20R | NR_015539.1 | 397-496 | CTAACAGAATTCCGTGTCACTCTGAATAGCACTCGAACTTCAGGCATTGGATTCCAGGCCTTGGACACAGTAGGAATCTGCTGTATCAATGAAGGATGTG | 80 | 82 | | PMO | 9630013A2C | NR_015539.1:396 | |
| 49 | Nfasc | NM_001160316.1 | 4936-5035 | CTCTTCGTATGCCTTATACAGCTCGGATCTAACCCTGCGGTTCCCAGATCCCTAGCCCTATTCTGCAAGTCTTGATGCCTCCAGTGATGTGTGTCTCGGG | 81 | 82 | | PMO | Nfasc | NM_001160316.1:4935 | |
| 50 | Ssh3 | NM_198113.2 | 1131-1230 | CTTCAGAAAAACAGAGTAAGTCACATCTTGAACATGGCCCGGGAGATTGACAACTTCTTCCCTGAGCGCTTCACCTATTACAATGTACGTGTCTGGGATG | 82 | 79 | | PMO | Ssh3 | NM_198113.2:1130 | |
| 51 | Pik3r3 | NM_181585.5 | 3091-3190 | TTCCGTGGCTGTGGTTTAAGCCTTCAGCTAAGTGAAGTTAGATAGAGGAGAGGGCAGCCATCTATTTATGGATTCACACTCATTTAAGAGTTCAGCTGCT | 80 | 78 | | PMO | Pik3r3 | NM_181585.5:3090 | |
| 52 | Enpp6 | NM_177304.3 | 923-1022 | GTGGAACACATGACCGTGTATGAGAAAGAATCAATACCCAACAGGTTCTATTACAAGAAAGGAAAATTTGTCTCTCCTTTGACCCTGGTGGCTGATGAAG | 83 | 80 | | PMO | Enpp6 | NM_177304.3:922 | |
| 53 | Tns3 | NM_001083587.1 | 4006-4105 | CGGCATTTCCTGATTGAGTGTACCCCAAAGGGAGTGCGGTTGAAAGGGTGCTCAAATGAACCATATTTTGGGAGCCTGACGGCTTTGGTGTGCCAGCATT | 82 | 81 | | PMO | Tns3 | NM_001083587.1:4005 | |
| 54 | Bmp4 | NM_007554.2 | 1501-1600 | AGACCCTAGTCAACTCTGTTAATTCTAGTATCCCTAAGGCCTGTTGTGTCCCCACTGAACTGAGTGCCATTTCCATGTTGTACCTGGATGAGTATGACAA | 79 | 82 | | PMO | Bmp4 | NM_007554.2:1500 | |
| 55 | Mycl1 | NM_008506.2 | 2111-2210 | AAACCTTTAAGATAGATTTGGATGGTAGGCCACACCCTTCCCTGCACGCTCAATGCTATGACTTTGAGAAAGGGCTTGGCCTCTATGTAGAGTCTTTGTC | 82 | 79 | | PMO | Mycl | NM_008506.2:2110 | |
| 56 | Cdv3 | NM_175833.2 | 1159-1258 | TAGGGAGGAGGTTTCAAAAAACCAGGCCCTTAAACTTCAGCTAGACAACCAGTATGCTGTGCTTGAGAATCAGAAGTACAGCCACACACAGTACAGCTAA | 85 | 86 | | PMO | Cdv3 | NM_175833.2:1158 | |
| 57 | Tmem163 | NM_028135.2 | 915-1014 | GACCTCTCAACTAGGCTGCTCCCTGAAGTGGATGATTTCCTGTTCAGCGTCTCCATTCTAAGCGGGATCCTTTGCAGCGTACTGGCTGTGCTGAAGTTTA | 82 | 81 | | PMO | Tmem163 | NM_028135.2:914 | |
| 58 | Rap2a | NM_029519.3 | 786-885 | GACCCATGCTGTTCTGCCTGTAACATACAATAGCATCCGAACATGGCTGCCCCGGACAGGATTCGTCTAGGCTCGTAGGCAACAGTAACCTTGTCTACTG | 83 | 82 | | PMO | Rap2a | NM_029519.3:785 | |
| 59 | Tmem2_tv2 | NM_001033759.2 | 398-497 | GCCATCGCGTAGACACCTGAGTAGACCGACGCCAGGAGGGGGAGGGGAAGTGACAGGGTATCATGTATGCCGCTGGTTCCAGGGGCCACTCCCCTGCTTT | 96 | 85 | T | PMO | Tmem2 | NM_001033759.2:397 | targets variant 2 only |
| 60 | Cnksr3 | NM_172546.2 | 2701-2800 | ATATTTATAAGTATGGCAGCGTCCGCCACAGAATGATGTGTGTGAGGCTCTTTGTAGCAGGGCTTTCTTTACCTACAGATCAGCTGTTACCCGCGCCCAG | 81 | 81 | | PMO | Cnksr3 | NM_172546.2:2700 | |
| 61 | Cyfip2 | NM_001252459.1 | 1137-1236 | TATGAGAATAAGATGTACCTGACGCCCAGTGAAAAGCACATGTTGCTCAAGGTGATGGGCTTTGGTCTCTATCTGATGGACGGAAATGTCAGTAACATTT | 82 | 80 | | PMO | Cyfip2 | NM_001252459.1:1136 | |
| 62 | Frmd4a | NM_001177843.1 | 3821-3920 | TTCCAAGTCACCTCACTGCCTCTCATTTGCCTTACCCAGATGCACTGTCACCTGCACCAGCTTCGGCCCTCAGCACTTTTCTTCTCCTGTCTCCACATCC | 82 | 82 | | PMO | Frmd4a | NM_001177843.1:3820 | |
| 63 | Slc12a2 | NM_009194.3 | 3746-3845 | CATAACTGTTCTGCACCATGGACTTCCTCTAAGGATGGCACTTCAGTGCCCAGTCAAAGTGACTGTATCGTCAGGAACACCACAGCATCGCAGTGGCAGA | 82 | 82 | | PMO | Slc12a2 | NM_009194.3:3745 | |
| 64 | Itpr2 | NM_010586.1 | 4366-4465 | GTGTGTGACAGAGTCAGTCATGAACATTGTGAGCGGCTTCTTCAACTCGCCATTTTCCGACAACAGTACCAGCCTCCAGACCCACCAGCCAGTCTTTATC | 82 | 82 | | PMO | Itpr2 | NM_010586.1:4365 | |
| 65 | Rnf122 | NM_175136.2 | 296-395 | TTCATTGATGCACCCATTCCAGTGGTGTAACGGGTGTTTCTGTGGCCTGGGACTGGTTAGTACCAACAAGTCCTGCTCAATGCCACCGATCAGTTTCCAA | 81 | 82 | | PMO | Rnf122 | NM_175136.2:295 | |
| 66 | Lims2 | NM_144862.3 | 869-968 | ACTGCTTTTCCTGTTCCACCTGCAACATGAAGCTCACTCTAAAGAACAAGTTTGTGGAGTTCGATATGAAGCCCGTGTGTAAGAGGTGCTATGAAAGGTT | 82 | 79 | | PMO | Lims2 | NM_144862.3:868 | |
| 67 | Samd4b | NM_175021.3 | 2403-2502 | AGCGAGAGGGTGGGCTGGCGCAGCTATACATTCTGATATTTTTCTACTCATCCTCTTAACTTTTGTTTAACATTGGCACACGCCTTGCTCACTCCCAGGC | 82 | 79 | | PMO | Samd4b | NM_175021.3:2402 | |
| 68 | Chn2 | NM_001163640.1 | 1511-1610 | ACCCTCCATGACATGCGGTACCAAAAGCTGATCGTGCAGATTTTAATAGAAAACGAAGATGTTCTGTTTTAATCCACCAGGGAAGTGAGCTGATAGAGCT | 81 | 81 | | PMO | Chn2 | NM_001163640.1:1510 | |
| 69 | Ppp2r3a | NM_001161362.3 | 2721-2820 | TTGTCCATAAACATTCCACAGTTCTATTTCCCTGAAGGACTCCCAGACACCTGCAGTAACCATGAACAGATCCTAAGCAGAATTGAAACTGCTTTCAAGG | 79 | 82 | | PMO | Ppp2r3a | NM_001161362.3:2720 | |
| 70 | Strn | NM_011500.2 | 1687-1786 | GCGGATCCAGAGCTGGAGCACCACCAATCCCAACGTCGATCCCTACGACGCTTACGATCCTTCAGTTCTAAGAGGCCCTCTGCTGGGCCACACGGACGCG | 83 | 82 | | PMO | Strn | NM_011500.2:1686 | |

| | | | | | | | | | | | |
|---|---|---|---|---|---|---|---|---|---|---|---|
| 71 | Glrb | NM_010298.5 | 705-804 | TTTCCCATGGACACACAACGCTGCAAAATGCAACTTGAGAGCTTTGGATATACAACCGATGATTTAAGATTCATCTGGCAGTCAGGAGATCCTGTTCAGT | 82 | 78 | | PMO | Glrb | NM_010298.5:704 | |
| 72 | Rras2 | NM_025846.2 | 1866-1965 | AAGTCTCACTGTTCACTGTTAACTGACAAAGAGACCATGGCTTTTGCTTGCTGGCTTGGTCCTTTTTACATCCACTGGTGGCCAATAATCAGAAAACCTT | 78 | 79 | | PMO | Rras2 | NM_025846.2:1865 | |
| 73 | Fmnl2 | NM_172409.2 | 959-1058 | ACGAGCCATCATGAATTACCAGTATGGCTTCAACATGGTCATGTCTCATCCACACGCCGTCAATGAGATTGCACTAAGTCTGAACAACAAAAATCCCAGA | 80 | 83 | | PMO | Fmnl2 | NM_172409.2:958 | |
| 74 | Sema5a | NM_009154.2 | 3821-3920 | CTCACTGGGAAGACCTATTCCAACGCCTACTTTACAGATCTCAACAATTATGATGAATACTAGCAGCTCTTATACTTTGGGCTCGTTGTAAACTCGCTGC | 80 | 80 | | PMO | Sema5a | NM_009154.2:3820 | |
| 75 | Fam3c | NM_138587.4 | 2601-2700 | GTCTTGTACTGGTTACACCGGCTCAACTCGGGAAGCAATAATCGACTGTTTTATGTTCACATCAGCCTTAGAAGTGGAACAGTGCTCACTTACTGTTGAC | 82 | 79 | | PMO | Fam3c | NM_138587.4:2600 | |
| 76 | Cdc37l1 | NM_025950.2 | 583-682 | TTTTGTCTGACCACCCACACCTCGTATGTGAAGAGACTGCTAAATATCTTATTTTATGGTGTTTCCACCTAGAAGCTGAGCAGAAAGGCGCTCTAATGGA | 79 | 81 | | PMO | Cdc37l1 | NM_025950.2:582 | |
| 77 | Fam73a | NM_001162375.1 | 877-976 | TAACATTCCTGTGACCACACCTGAGAACTTATACCTGATGGGTATGGAATTATTTGAAGAGGCGTTACGACGATGGGAACAAGCTCTGACTTTTCGAAGT | 82 | 79 | | PMO | Miga1 | NM_001162375.1:876 | |
| 78 | Elovl6 | NM_130450.2 | 4196-4295 | AAGACTGGAGGGGAAATCCATATTCATTGCTGCAGGGGAAAGCAGGTTTATTAATCCTCCCGCTGTCTTTATGAGACCGATTGACCAATGTAGCTCAGGC | 80 | 79 | | PMO | Elovl6 | NM_130450.2:4195 | |
| 79 | Atrn | NM_009730.2 | 1376-1475 | TAAGGATCAGTATGCAGTGGTTGGACACTCAGCACACATTGTTACACTGGCATCTGGCCGTGTGGTCATGTTGGTCATCTTCGGTCATTGCCCACTCTAT | 82 | 79 | | PMO | Atrn | NM_009730.2:1375 | |
| 80 | Tmem2_tv1 | NM_031997.4 | 398-497 | GCCATCGCGTAGACACCTGAGTAGACCGACGCCAGGAGGGGGAGGGGAAGAACGGAAAGTTAAGGTTAGACGAGAACTCTGAGCCGTCACCTGTTCTCTG | 96 | 83 | T | PMO | Tmem2 | NM_031997.4:397 | targets variant 1 only |
| 81 | Gjb1 | NM_008124.2 | 114-213 | GTCTGTCATCTTCATCTTCAGAATCATGGTGCTGGTGGTGGCTGCTGAGAGCGTGTGGGGGGATGAGAAGTCCTCTTTCATCTGTAACACCCTCCAGCCG | 83 | 82 | | MO | Gjb1 | NM_008124.2:113 | |
| 82 | Ndrg1 | NM_008681.2 | 2326-2425 | ATTGATTTGAAACTTGATAGCTGAATGTCTCTTAACTGGCCCATCGCTGGTTCTTCAGAGAGAACCGTATGTTCTAGTGGGGGAAGGACTGGTTTTGGGT | 78 | 82 | | MO | Ndrg1 | NM_008681.2:2325 | |
| 83 | Ppp1r14a | NM_026731.3 | 274-373 | GAGGAAAGATGCCGGAAAATCCAGGGACTTCTGGAGGCTTGTGCAAATCCCACAGAGGACTTCGTCCAGGAGCTACTGGCCAAGCTTCGGGGCCTGCACA | 84 | 86 | | MO | Ppp1r14a | NM_026731.3:273 | |
| 84 | Adssl1 | NM_007421.2 | 1671-1770 | TAAAATGAGCCTAAGTGCTCATAACAGAGGCCTTCACTGACTGATTGTCACATGGTATGTGTTCAGTTGGAGCCATGTCATAATAAACATAGCCAGGGGT | 83 | 81 | | MO | Adssl1 | NM_007421.2:1670 | |
| 85 | Aspa | NM_023113.5 | 175-274 | CCTATTAAGAAGATTGCCATCTTTGGAGGGACTCATGGAAATGAACTGACCGGAGTGTTTCTAGTTACTCACTGGCTAAGGAATGGCACTGAAGTTCACA | 82 | 81 | | MO | Aspa | NM_023113.5:174 | |
| 86 | Acy3 | NM_027857.3 | 1159-1258 | CTTTGAGCCACTGAGGCCTGGTGAACCCATCTTCAAGCTTTTCAGCGGAGAAGACGTACTGTATGAGGGGGACTCCATTGTGTACCCTGTGTTCATTAAT | 81 | 81 | | MO | Acy3 | NM_027857.3:1158 | |
| 87 | Trp53inp2 | NM_178111.3 | 2091-2190 | CTGGGTAGAGGAGGCCAGGAGCCTCACTGTCCCAATTCCCAAATTTTTCTTACAGAGATTCAGGGCCTTCAAGAGTCTTCTTAATGTTCTCCCACAGGGC | 81 | 80 | | MO | Trp53inp2 | NM_178111.3:2090 | |
| 88 | Pla2g16 | NM_139269.2 | 1021-1120 | GAACTTGTTTCAACGTGGCGCTTTGAGACTCAGGCGCTGGAGTTTATGGATATTCAGTGGTGTGAGAAGGACAAATCCTGGAGTGTTGGGGCTTTAGCTC | 81 | 82 | | MO | Pla2g16 | NM_139269.2:1020 | |
| 89 | Efhd1 | NM_028889.2 | 1582-1681 | CAACAGTTCCAGAGAACTGAAAGAAATGTATTCTCGCCCATGAGACAGATTTTTGTATCTTGAGTTTCTTGGGACTTCTCACAAATACCTGGGGAGGGGG | 82 | 79 | | MO | Efhd1 | NM_028889.2:1581 | |
| 90 | Itgb4 | NM_001005608.2 | 3356-3455 | TGCCACCCTTGGTCGCCGTCTGGTAAACATCACCATTATCAAGGAACAAGCTAGTGGGGTAGTGTCCTTCGAGCAGCCTGAATACTCGGTGAGTCGTGGA | 82 | 83 | | MO | Itgb4 | NM_001005608.2:3355 | |
| 91 | Hapln2 | NM_022031.2 | 1827-1926 | CTCTACCTCCTAAGTTCTGGGATCACATCTGGCTAAAGTCTACAATGAAAGACGGTAAATTCCGTGTGGACTGAGAACTCAGCAGCTAAGCTAACTTGAT | 81 | 82 | | MO | Hapln2 | NM_022031.2:1826 | |
| 92 | mbp | NM_010777.3 | 762-861 | TGCCAGGCATGGCTTCCTCCCAAGGCACAGAGACACGGGCATCCTTGACTCCATCGGGCGCTTCTTTAGCGGTGACAGGGGTGCGCCCAAGCGGGGCTCT | 86 | 85 | | MO | Mbp | NM_010777.3:761 | |
| 93 | Hcn2 | NM_008226.2 | 1901-2000 | GCGTGCCTTTGAGACTGTGGCTATTGACCGGCTAGATCGCATAGGCAAGAAGAACTCCATCTTGCTGCACAAGGTTCAGCATGATCTCAGCTCAGGTGTG | 82 | 82 | | MO | Hcn2 | NM_008226.2:1900 | |
| 94 | Nmra1 | NM_026393.1 | 551-650 | CAAAGGGGAGGTGGAGGAATACTTCCGAGACATCGGTGTTCCCATGACCAGTGTGCGGCTGCCTTGCTATTTCGAGAATCTCCTTTCCTATTTCCTGCCC | 82 | 78 | | MO | Nmral1 | NM_026393.1:550 | |
| 95 | Cdc42ep2 | NM_026772.2 | 1369-1468 | CTTTGTGTTTGGGTCTGGGAGTCCAGAAATCGTGACATCTCCCGCCTTGTGTCCTGTCCCAACCCTTCCTCACATGACCAGGGAATTCTGGTCGCAATAA | 80 | 82 | | MO | Cdc42ep2 | NM_026772.2:1368 | |
| 96 | Mal | NM_001171187.1 | 686-785 | TTTTACCTCAGTGCCTCAGTTCTGGAAGCCCTGGCCACCATCTCAATGTTTGATGGCTTTACTTACAAGCATTACCATGAAAACATCGCCGCAGTGGTGT | 82 | 81 | | MO | Mal | NM_001171187.1:685 | |
| 97 | Mog | NM_010814.2 | 751-850 | AAAACTTCGTGCAGAAGTAGAGAATCTCCATCGGACTTTTGATCCTCACTTCCTGAGGGTGCCCTGCTGGAAGATAACACTGTTTGTTATTGTGCCTGTT | 79 | 82 | | MO | Mog | NM_010814.2:750 | |
| 98 | Slco3a1 | NM_001038643.1 | 856-955 | GGTATTTGGACCAGCCTGTGGATTTATTCTGGGCTCTTTCTGTACCAAGATCTATGTGGATGCGGTCTTCATTGACACAAGTAACCTGGACATCACTCCA | 79 | 83 | | MO | Slco3a1 | NM_001038643.1:855 | |
| 99 | Apod | XM_003689301.1 | 351-450 | CATCTTGGGAAATGCCCGTCTCCTCCTGTGCAAGAGAATTTTGACGTGAAAAAGTATCTTGGAAGATGGTACGAAATTGAGAAGATCCCAGCGAGCTTTG | 80 | 80 | | MO | Apod | XM_003689301.1:350 | |
| 100 | Gsn | NM_146120.3 | 625-724 | GTACCTGTGTCCTGGGACAGTTTCAACAATGGCGACTGCTTCATTCTGGACCTGGGAAACAATATCTATCAGTGGTGTGGCTCTGGCAGCAACAAATTTG | 83 | 83 | | MO | Gsn | NM_146120.3:624 | |
| 101 | Pdlim2 | NM_145978.1 | 1366-1465 | CCATGTTTGACACGAAAGCGTATGGGGTTGGGCATGGATAGAAGCATCTAGAAGGGAATGGTGGGCCTGAGGTAAATGATATTCATGGTGTGAAGTTTCT | 82 | 82 | | MO | Pdlim2 | NM_145978.1:1365 | |
| 102 | Prr18 | NR_028280.1 | 2019-2118 | TCTCCAATGGCACAGCTGGAATTGTTTTCAGAAGTTGGATCAAGGTGGGGACAGGAGTCAGGGTTGTCGTCTGCTGGGATTGCAGTTTTTATTTGAGGGC | 82 | 80 | | MO | Prr18 | NR_028280.1:2018 | |
| 103 | Inf2 | NM_198411.2 | 677-776 | TCGGACAGTGACAACGTGCCCTATGTGGTCACCCTGCTCAGTGTGATCAACGCCATCATTCTGGGTCCCGAGGACCTCCGCTCTCGTGCCCAGCTGCGGA | 82 | 83 | | MO | Inf2 | NM_198411.2:676 | |
| 104 | Tppp3 | NM_026481.3 | 915-1014 | TCCCAGGCATGGCTGACTCTTGCCTGCTTTTGCCTCATATTTAAGCTGCTGCTCTGGCCAAGTGCCTAATTCTTACCCAGACCTCAATAAAGATACCTTT | 83 | 81 | | MO | Tppp3 | NM_026481.3:914 | |
| 105 | Tbc1d9b | NM_029745.2 | 351-450 | AAAACAACTTGCTTCAGACACTGTCCATCTTCGACAACGAGGAGGACATCACTACCTTCGTCAAGGGCAAGATACACGGCATCATCGCAGAAGAGAACAA | 82 | 82 | | MO | Tbc1d9b | NM_029745.2:350 | |
| 106 | Nol3 | NM_030152.4 | 917-1016 | AGACACCTACTGCCAAATGAATAAACTCAGGAGGGTCAGTCTGTTCCTGACTCTCCACAAGTCTGGCTCTTTGCTCAAGAACTGAGGGTGAGCGTGGGTC | 82 | 82 | | MO | Nol3 | NM_030152.4:916 | |
| 107 | Cenpb | NM_007682.2 | 1248-1347 | CCTGGCAGGCAGTGGAGCCCTCGGACATAGCAACTTGCTTTCGTGAGGCGGGTTTTGGAGGTGGCCTTAATGCCACTATCACCACTTCCTTCAAGAGTGA | 90 | 81 | | MO | Cenpb | NM_007682.2:1247 | |
| 108 | Slc45a3 | NM_145977.1 | 2331-2430 | TGCAGGGATTGAAATGTGGTCTGTTCAGGTGTCCTGTATCAAGGCTCCAGGTTAACAGCTAACCTCCTGCTTAAGTCACCAGAGGGAGGATTTGGGGGAC | 80 | 82 | | MO | Slc45a3 | NM_145977.1:2330 | |
| 109 | Carns1 | NM_134148.2 | 3729-3828 | TCTTCATTTATAGGCAGCACTCAGACTCCTAAACACCTGCCTCCCATACACCACACACGATCTCTACCAGCTTTGTTGCTTTGGCCTCTGGCATTCCTAG | 83 | 81 | | MO | Carns1 | NM_134148.2:3728 | |
| 110 | Opalin | NM_153520.1 | 765-864 | GTATGTCCTGTTTCTGAATCGAGAGAAGTCTGCCTTTGGGAAACGGCTTGGCTCCCGAGTGTCCTGCCTTCTCCTCTACCTGCTCCAAATGTTATCATTT | 80 | 79 | | MO | Opalin | NM_153520.1:764 | |
| 111 | Arsg | NM_028710.3 | 2897-2996 | TGTTGCCTTTGAGATGACTTGAGCAAGCTAAACCAAAGTGAGGTTTTCCCAGGCCTTGGACCCTTCTCTCCTGTAAATGTCAAGTGGATGTATTATAAAG | 80 | 79 | | MO | Arsg | NM_028710.3:2896 | |
| 112 | Rftn1 | NM_181397.2 | 1559-1658 | TCCACCAAGCAGATTGTGTTTCTTCAGAGACCATGTCTACCTCAGAAAACCAAGAAGAGGGAATCTAAGTTCCAGTGGCGATTCTCCCGGAACGAGATCC | 81 | 82 | | MO | Rftn1 | NM_181397.2:1558 | |
| 113 | Adap1 | NM_172723.4 | 429-528 | CCGTGCCAAGTATGAGAGGCAAGAATTCGTTCATGTGGAGAAGCAGGAACCATACTCTACAGGCTACCGTGAGGGCCTTCTCTGGAAGCGTGGGCGGGAC | 82 | 83 | | MO | Adap1 | NM_172723.4:428 | |
| 114 | Plekhb1 | NM_001163184.1 | 1617-1716 | GTGCCTGTCTATAGGTAGCAAGCAATCTCTCCATTTGATCCTGTTCCAAACTATTCCATCATCCTGTCCCAATTGCCAATAGAAAAGCTAAAATCCCTGG | 78 | 79 | | MO | Plekhb1 | NM_001163184.1:1616 | |
| 115 | Trf | NM_133977.2 | 1941-2040 | AGGAGACTTTATTTGGGGGAAGTGACTGCACCGGCAATTTCTGTTTGTTCAAGTCCACCACCAAGGACCTTCTGTTCAGGGATGACACCAAATGTTTCGT | 79 | 83 | | MO | Trf | NM_133977.2:1940 | |
| 116 | Insc | NM_173767.3 | 1475-1574 | AAACATGTCTGTGCTGGAGCAGTGTGGCTCTGACATCATTCAGGAGAACGGTGTCCAGCTCATCATGGGCATGCTGTCTGAGAAGCCAAGGTCTGGGACA | 83 | 82 | | MO | Insc | NM_173767.3:1474 | |
| 117 | Cryab | NM_009964.2 | 919-1018 | ACTTCATCCCTGTCATCTGATGGAGTCCTCACTGTGAATGGACCAAGGAAACAGGTGTCTGGCCCTGAGCGCACCATTCCCATCACCCGTGAAGAGAAGC | 85 | 85 | | MO | Cryab | NM_009964.2:918 | |
| 118 | Kif5a | NM_008447.4 | 879-978 | ATTTACCTGGATAAGATCCGTGACCTTTTGGATGTGACCAAGACGAACCTGTCCGTGCATGAGGACAAAAACCGGGTGCCGTTTGTCAAGGGTTGTACCG | 81 | 81 | | MO | Kif5a | NM_008447.4:878 | |
| 119 | Trak2 | NM_172406.3 | 2833-2932 | GTCTCCGAGAATTTCTTGGCTTCCCGACCAGCTGAAACATTCCTGCAAGAAATGTATGGCTTGAGACCTTCAAGGGCCCCTCCTGATGTTGGACAGCTGA | 82 | 88 | | MO | Trak2 | NM_172406.3:2832 | |
| 120 | Cldn11 | NM_008770.3 | 1175-1274 | GTCTTGGGAACATTGCTGACTTGGGTGTGGATTGGGAAAGAAATAAAAGATGCTTTTCAAAGGGTTACCAACGACAGTGGAAGCCTTATAGAGACAGCTC | 83 | 81 | | MO | Cldn11 | NM_008770.3:1174 | |
| 121 | gapdh | NM_001001303.1 | 891-990 | AGGTTGTCTCCTGCGACTTCAACAGCAACTCCCACTCTTCCACCTTCGATGCCGGGGCTGGCATTGCTCTCAATGACAACTTTGTCAAGCTCATTTCCTG | 81 | 79 | HK;X | HK | Gapdh | NM_001001303.1:890 | targets multiple predicted and non-coding RNAs @ 100% |
| 122 | rpl19 | NM_009078.1 | 21-120 | GAAGAGGCTTGCCTCTAGTGTCCTCCGCTGCGGGAAAAAGAAGGTCTGGTTGGATCCCAATGAGACCAATGAAATCGCCAATGCCAACTCCCGTCAGCAG | 83 | 80 | HK | HK | Rpl19 | NM_009078.1:20 | |
| 123 | actb | NM_007393.1 | 816-915 | CAGGTCATCACTATTGGCAACGAGCGGTTCCGATGCCCTGAGGCTCTTTTCCAGCCTTCCTTCTTGGGTATGGAATCCTGTGGCATCCATGAAACTACAT | 81 | 79 | HK;X | HK | Actb | NM_007393.1:815 | also targets Lrrc58 (NM_177093) @97% |
| 124 | pgk1 | NM_008828.2 | 37-136 | CCGGCATTCTGCACGCTTCAAAAGCGCACGTCTGCCGCGCTGTTCTCCTCTTCCTCATCTCCGGGCCTTTCGACCTCACGGTGTTGCCAAAATGTCGCTT | 83 | 80 | HK | HK | Pgk1 | NM_008828.2:36 | |
| 125 | b2m | NM_009735.3 | 178-277 | CTGAACTGCTACGTAACACAGTTCCACCCGCCTCACATTGAAATCCAAATGCTGAAGAACGGGAAAAAAATTCCTAAAGTAGAGATGTCAGATATGTCCT | 82 | 81 | HK | HK | B2m | NM_009735.3:177 | |
